# Comprehensive single cell transcriptomics analysis of murine osteosarcoma uncovers *Skp2* function in metastasis, genomic instability and immune activation and reveals additional target pathways

**DOI:** 10.1101/2024.06.04.597347

**Authors:** Alexander Ferrena, Ranxin Zhang, Jichuan Wang, Xiang Yu Zheng, Barlas Göker, Hasibagan Borjihan, Sung-Suk Chae, Yungtai Lo, Hongling Zhao, Edward Schwartz, David Loeb, Rui Yang, David Geller, Deyou Zheng, Bang Hoang

## Abstract

Osteosarcoma (OS) is the most common primary pediatric bone malignancy. One promising new therapeutic target is *SKP2*, encoding a substrate recognition factor of the SCF E3 ubiquitin ligase responsible for ubiquitination and proteasome degradation of substrate p27, thus driving cellular proliferation. We have shown previously that knockout of *Skp2* in an immunocompetent transgenic mouse model of OS improved survival, drove apoptosis, and induced tumor inflammation. Here, we applied single-cell RNA-sequencing (scRNA-seq) to study primary OS tumors derived from Osx-Cre driven conditional knockout of *Rb1* and *Trp53*. We showed that murine OS models recapitulate the tumor heterogeneity and microenvironment complexity observed in patient tumors. We further compared this model with OS models with functional disruption of *Skp2*: one with *Skp2* knockout and the other with the Skp2-p27 interaction disrupted (resulting in p27 overexpression). We found reduction of T cell exhaustion and upregulation of interferon activation, along with evidence of replicative and endoplasmic reticulum-related stress in the *Skp2* disruption models, and showed that interferon induction was correlated with improved survival in OS patients. Additionally, our scRNA-seq analysis uncovered decreased activities of metastasis-related gene signatures in the *Skp2*-disrupted OS, which we validated by observation of a strong reduction in lung metastasis in the *Skp2* knockout mice. Finally, we report several potential mechanisms of escape from targeting *Skp2* in OS, including upregulation of *Myc* targets, DNA copy number amplification and overexpression of alternative E3 ligase genes, and potential alternative lineage activation. These mechanistic insights into OS tumor biology and *Skp2* function suggest novel targets for new, synergistic therapies, while the data and our comprehensive analysis may serve as a public resource for further big data-driven OS research.

## INTRODUCTION

Osteosarcoma (OS) is a primary bone malignancy defined by spindle cell morphology and abnormal deposition of bone matrix^1,2^. OS is the most frequent pediatric bone cancer with annual US incidence of 3.1 cases per million^3^. Disease progression usually involves metastasis to the lungs^4^. For cases that are non-metastatic at diagnosis, 5-year survival is 70% while for metastatic cases at presentation, comprising 15 to 25% of incidence, the rate is 30%^4,5^. Standard of care treatment for OS involves cytotoxic chemotherapy and resection, a regimen unchanged since 1986^6^. Genetically, OS is complex and involves many DNA copy number alterations, but the two most frequent mutations are loss of the tumor suppressors *TP53* and *RB1*^7–9^. Histologically, several OS subtypes have been described, including osteoblastic OS, chondroblastic OS, and fibroblastic OS, but the molecular distinctions among them and functional implications are unknown, and moreover clinical management and outcomes are similar between these subtypes^10^.

S-phase Kinase Associated Protein 2 (*SKP2*) codes for a substrate recognition factor of the Skp1-Cullin1-F-box (SCF) E3 ubiquitin ligase complex. One key target for ubiquitination and proteasome degradation by the SCF^SKP2^ complex is the cyclin-dependent kinase inhibitors p27, allowing G1-S phase transition during cell cycle progression^11,12^. *Skp2* knockout was shown to be synthetic-lethal in mouse models of pituitary and prostate cancers in the context of *Rb1* and *Trp53* loss, permanently blocking tumorigenesis in a p27-mediated manner^13^. Mechanistically, *Skp2* deletion resulted in hyperactivation of the pleiotropic transcription factor *E2f1*, resulting in both increasing signatures of proliferation but also, critically, in malignant cell apoptosis^14^. In transgenic mouse models of OS driven by conditional ablation of *Rb1* and *Trp53* in the bone-forming Osx (*Sp7*) lineage, we showed that genetic and pharmacologic inhibition of *Skp2*’s interaction with p27 improved survival, decreased tumor size, and reduced stemness; however, while delayed, tumorigenesis and disease progression still occurred, suggesting a resistance or escape mechanism^15,16^. Furthermore, we showed that knockout of *Skp2* resulted in even greater survival improvement and tumor apoptosis in a manner related to *E2f1*^17^. Surprisingly, in addition to *E2f1* activity, we also detected increased inflammation in the form of immune infiltration and interferon expression in *Skp2* knockout OS tumors, with the increases associated with improved prognosis in OS patients, suggesting a role of *Skp2* in immunosuppression in OS^18^. However, in a distinct cancer context, *Myc*-driven Burkitt lymphomagenesis, *Skp2* knockout had only a modest improvement in survival^19^. The mechanism employed by OS to escape from *Skp2* disruption, along with the regulatory roles of *Skp2* in the tumor microenvironment, remain unknown.

As previously reported, conditional ablation of *Rb1* and *Trp53* from the bone-forming Osx (Sp7) lineage (“Double Knockout”, “DKO”: Osx-Cre;Rb1^lox/lox^;p53^lox/lox^) forms tumors that strongly resemble OS patient tumor histology and disease progression^15^. We compared this *Skp2*-intact baseline OS model against two models of *Skp2* genetic modulation. The first involved disruption of *Skp2*’s interaction with p27 via a mutation preventing p27 phosphorylation and *Skp2* binding, resulting in a “p27-high” phenotype (“Double Knockout Alanine-Alanine”, “DKOAA”: Osx-Cre;Rb1^lox/lox^;p53^lox/lox^; p27^T871A/T187A^). The second involved a germline *Skp2* knockout (“Triple Knockout”, “TKO”: Osx-Cre;Rb1^lox/lox^;p53^lox/lox^; Skp2^-/-^). In the DKOAA and TKO models, we observed improved survival, delayed tumorigenesis, reduced tumor size, increased tumor inflammation, and induction of apoptosis, relative to DKO^16–18^.

Together these previous studies suggest that SKP2 disruption may have multifactorial roles, resulting in delayed but not blocked tumor formation eventually. How the different roles are executed and balanced in tumors can be addressed better using single cell transcriptomics (scRNA-seq) analysis to resolve cellular heterogeneity. Therefore, we applied scRNA-seq to compare the transcriptomes of the three transgenic murine OS tumor models. With these new data, we first investigated the mechanisms underlying the positive prompts of SKP2 disruption, including reduced metastasis, promoting anti-tumor microenvironment, and improved survival. We then studied the potential mechanisms used by OS to escape *Skp2* function loss. With the first reported scRNA-seq dataset of an animal model in OS, we also compared it to the scRNA-seq datasets from patient OS and validated that the murine OS models faithfully recapitulates the complex microenvironment and transcriptomic diversity of human OS. Taken together, our data are both valuable for addressing the therapeutic utility of SKP2 disruption and for advance further OS research in general.

## RESULTS

### A complex tumor microenvironment is present in the transgenic murine OS tumors

Ten tumors obtained from our three mouse models of OS (4 TKO, 3 DKOAA, 3 DKO) were submitted for scRNA-seq using the 10X Genomics platform [**Fig 1A, Supplementary Table 1**]. After strict quality control of the data, cells from all samples were integrated, clustered and annotated. In all samples, we detected malignant cells along with diverse immune cells (macrophages, osteoclasts, T cells, B cells, and neutrophils), and non-malignant stromal cells (endothelial cells and cancer-associated fibroblasts) [**Fig 1B, Supplementary Figure 1**]. This assortment of cell types closely matches the observations in two recent scRNA-seq studies of human patient OS tumors^20,21^. Cell types were annotated by the top markers computed from the data (i.e., data-derived markers) and additionally canonical markers for each cell type [**Fig 1C**]. Interestingly, malignant cells only expressed osteoblast lineage factor Osterix (*Sp7*) sparsely and were also positive for chondroblastic lineage factor *Sox9*, indicating lineage infidelity may be a feature of advanced OS tumors. While all the cell types were present in all models, transcriptomic distinctions in some cell types existed even after batch correction and data integration, such as malignant cells, indicating *Skp2’*s roles in OS development [**Fig 1D**].

**Figure 1:**
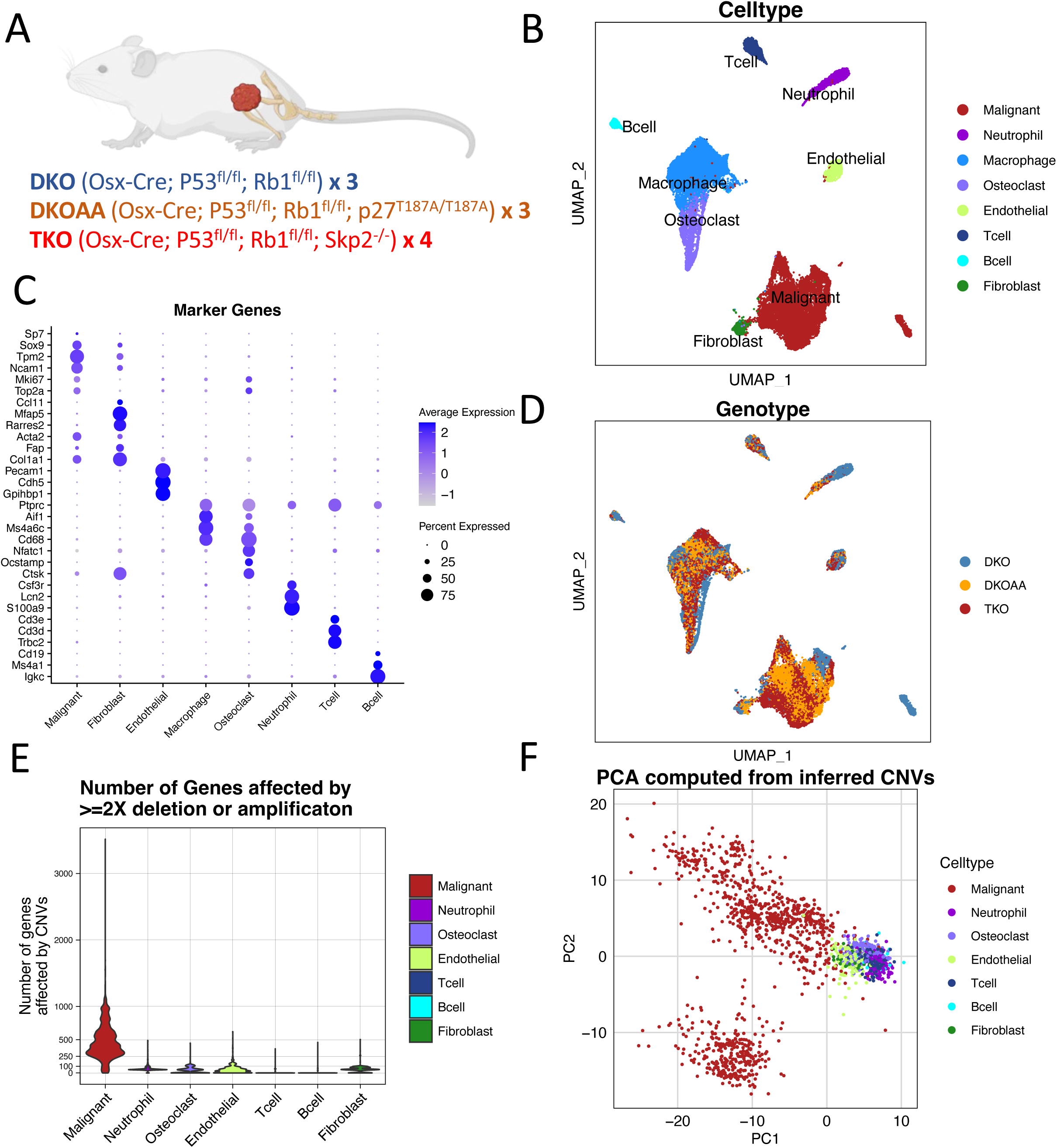
Complex microenvironment in transgenic murine OS tumors. A: Mouse models for scRNA-seq analysis. Numbers indicate the mice included, with one tumor isolated from each mouse for scRNA-seq. B: UMAP of integrated cells colored by cell types. C: Canonical and data-driven markers of cell types shown as bubble plot. D: UMAP colored by genotype. E: Violin plot of number of genes affected by “extreme CNVs” (deep 2x deletions, or 2x> amplification) in individual cell types. F: Principal Component Analysis (PCA) plot of all cells using their CNV profiles inferred from scRNA-seq data.

While most of the cell types could be annotated by their distinct transcriptomic profiles and marker genes [**Fig 1C, Supplementary Figure 1, Supplementary Tables 2-4**], the classification of malignant OS cells versus non-malignant mesenchymal stromal cells, especially fibroblasts, was less clear, likely reflecting the mesenchymal cell origin of OS. To resolve this, we called sequence mutations from the scRNA-seq data, considering that the malignant cells should have more mutations compared to their non-malignant counterparts. We started with single nucleotide variations and small indels using Souporcell, which clusters cells by their mutation profiles^22^. For most samples, such as DKO_1, DKOAA_1, and TKO_1, the analysis revealed a clear difference between suspected malignant OS cell clusters and non-malignant stromal cells [**Supplementary Figure 2**]. For some tumor samples, such as DKOAA_2, a complex multi-clonal malignant population appeared to be present, supported by two transcriptomically distinct clusters. For other samples, such as DKO_2, the separation of malignant and non-malignant cell types was more ambiguous and only possible after examining the integration co-clustering with cells from all samples.

OS tumors display high levels of genomic instability and high frequency of large-scale copy number variations (CNVs)^23^. We next turned to CNV analysis with the expectation that malignant cells would have more CNVs than stromal cells. For this, we used inferCNV^24^, which has been shown to identify CNVs reliably from scRNA-seq data at a level matching to the results called from whole exome sequencing of paired synovial sarcoma tumors^25,26^. Indeed, our inferCNV analysis detected clear differences between malignant and non-malignant cells, especially when considering the data for (2x) deletions of both gene copies or 2x or more amplifications (referred as “extreme CNVs”) [**Fig 1E, Supplementary Figure 3**]. To further support our identification of malignant cells, we performed principal component analysis (PCA) using the inferred CNV profiles. The results showed high complexity and diversity of CNVs among malignant cells, separating them from non-malignant cells that had significantly fewer CNVs regardless of cell type [**Fig 1F**].

In short, this comprehensive, integrative cell annotation approach enables us to reliably distinguish malignant cell clusters and all other cell types, thus allowing us to proceed with the downstream analysis with greater confidence. Additionally, this analysis revealed that *Rb1* and *Trp53* ablation in the Osterix (*Sp7*) lineage produces tumors that strongly resemble human OS in terms of microenvironment cell makeup and genetic instability (further discussion below).

### *Skp2* knockout reduces molecular signatures of invasiveness and incidence of lung metastasis in OS

In OS patients, disease progression typically manifests as aggressive metastasis to the lung, and if unresponsive to salvage treatment, results in death^4^. We previously reported that TKO and DKOAA mice both display significantly improved survival relative to DKO^17^. Additionally, knockdown of *SKP2* expression or pharmacologic *SKP2* inhibition in xenograft tumors derived from patient OS cell lines led to reduced metastasis in SCID mice^27^.

To study the effect of *Skp2* functional disruption on OS malignant cells, we performed differential expression analysis between TKO, DKOAA and DKO cells and uncovered many pathways significantly enriched among the differentially expressed genes (DEGs) [**Fig 2A, 2B; Supplementary Tables 5 and 6**]. Among the down-regulated pathways in both TKO and DKOAA malignant cells, Epithelial-Mesenchymal Transition (EMT) stands out (**Fig 2A and 2B bottom**).

**Figure 2:**
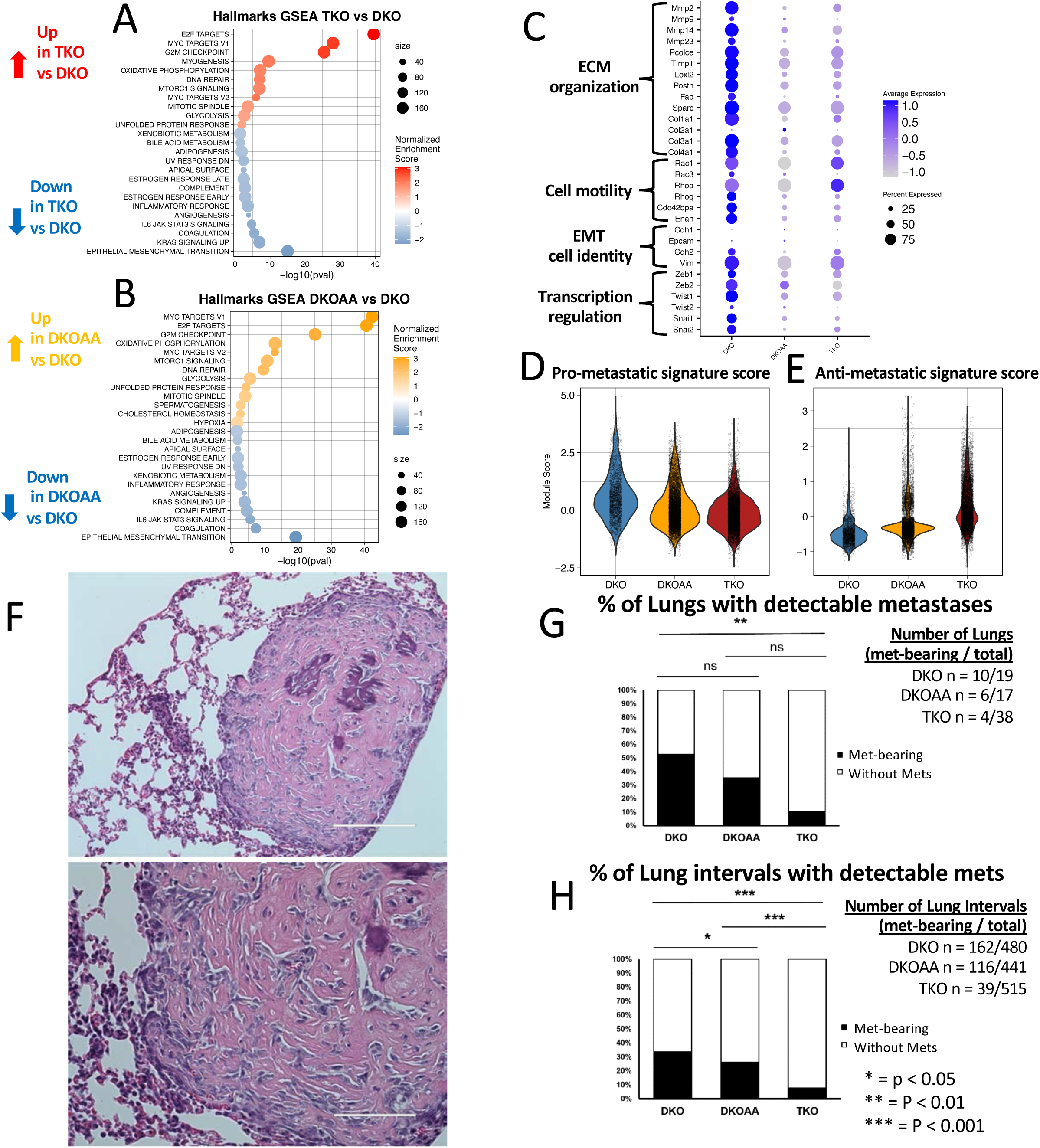
*SKP2* Knockout reducing OS lung metastasis and genes related to invasiveness. A: Dotplot of Hallmarks gene sets significantly more active in TKO relative to DKO in malignant cells. B: Dotplot of Hallmarks gene sets significantly more active in DKOAA relative to DKO in malignant cells. C: Dotplot showing aggregated single cell expression differences among OS models for genes in EMT related phenotypes including Extracellular Matrix (ECM) organization, cell motility, cell identity, and transcriptional regulation. D,E: Signature gene signatures scores of pro- and anti-metastatic OS genes^29^. F: Representative H&E image of lung metastasis tumor from TKO at 200 μm (top) and 100 μm (bottom) scales. G: Quantification and comparison of tumor-bearing lungs. H: Quantification and comparison of tumor-bearing serially sectioned lung intervals.

EMT has been linked to metastasis in many cancers including OS, and involves dramatic functional and cellular morphological changes. These include gain of cell motility and capacity for modification of the extracellular matrix, key features related to local invasiveness and distal metastasis^28^. Many genes related to extracellular matrix (ECM) organization and cell motility along with related transcription factors were downregulated in TKO [**Fig 2C**]. However, markers of epithelial cell identity were not strongly expressed in any tumor, indicating the difference is driven by functional phenotypes and that malignant OS cells did not activate the epithelial programs. Accordingly, we observed significant downregulation of gene sets related to ECM organization and cell motility in both TKO and DKOAA relative to DKO malignant cells [**Supplementary Figure 4].**

This expression difference in EMT-related genes suggests that TKO and DKOAA tumors may have reduced capacity for invasiveness and metastasis relative to DKO. To further investigate this, we used recently reported gene signatures to calculate pro- and anti-metastasis signature scores^29^. We found the highest pro-metastasis score in DKO tumors, an intermediate score in DKOAA, and the lowest pro-metastasis score in TKO [**Fig 2D**]. Conversely, we observed the opposite trends for the anti-metastasis scores (all P < 0.01, Wilcoxon test). [**Fig 2E**].

We next directly tested the incidence of lung metastasis in the three OS models. Using analysis of H&E stained FFPE lung samples, we were able to easily distinguish OS lung metastases from normal lung tissue [**Fig 2F**]. Quantification of H&E images from lungs revealed that the percent of lung metastases was significantly reduced in TKO compared to DKO (p < 0.05; Fisher exact test, **Fig 2G**). DKOAA tumors had a lower but non-significant reduction in metastasis-bearing lungs. To better quantify metastasis extent within each lung, we also analyzed serially sectioned lung intervals, and detected a significant reduction of metastasis-bearing lung intervals in both TKO and DKOAA relative to DKO, as well as a significant reduction in TKO relative to DKOAA [**Fig 2H**].

Taken together, our scRNA-seq analysis and *in vivo* study both indicate that *Skp2* knockout reduces the incidence of lung metastasis in OS. This suggests that *Skp2* disruption in OS has anti-tumor benefit by downregulation of EMT and metastasis-related gene expression.

### Functional disruption of SKP2 induces immune activation in the form of increased interferon pathway activity and reduction of T cell exhaustion

Our prior work on *SKP2* in OS revealed increased inflammation in TKO relative to DKO tumors, alongside evidence of increased infiltration of macrophages, T cells, B cells, and endothelial cells, as well as upregulation of interferon gamma response^18^. Additionally, we showed that TKO and DKOAA mice have improved survival compared to DKO, with tumors from these models showing increased tumor apoptosis and reduced stemness compared to DKO tumors^16,17^. We sought to expand on these findings, which were from bulk tumors, using scRNA-seq data to determine if the effects are cell type specific.

We thus performed differential gene expression analysis across TKO, DKOAA and DKO and identified DEGs that were significantly enriched in many pathways in individual cell types [**Fig 3A, Supplementary Figures 5-6**]. Notably, in all microenvironment cell clusters, interferon response pathways were significantly upregulated in TKO relative to DKO, with the strongest upregulation of both type 1 and type 2 interferon response observed in T cells [**Fig 3B**]. In DKOAA, interferon response pathways were also upregulated, but significant upregulation was restricted to T cells and osteoclasts [**Supplementary Figure 7A**]. Upregulated interferon-response genes were numerous and included transcription factors like *Irf5* and *Irf7*; antiviral genes such as *Oas2*; stress response regulatory genes such as *Eif2ak*; antigen presentation genes such as *H2-Q7*; tetratricopeptide-containing interferon-induced genes such as *Ifit3*; and pro-inflammatory cytokines such as *Cxcl10* [**Supplementary Figure 7**]. Interestingly, among upregulated genes in the interferon response pathways were E3 ubiquitin ligase genes such as *Trim21*, *Trim25* and *Trim26*. *Trim21* that have previously been demonstrated to bind with *SKP2*, causing degradation of p27^30^. We also found upregulation of *Stat1* in T cells [**Supplementary Table 5**].

**Figure 3:**
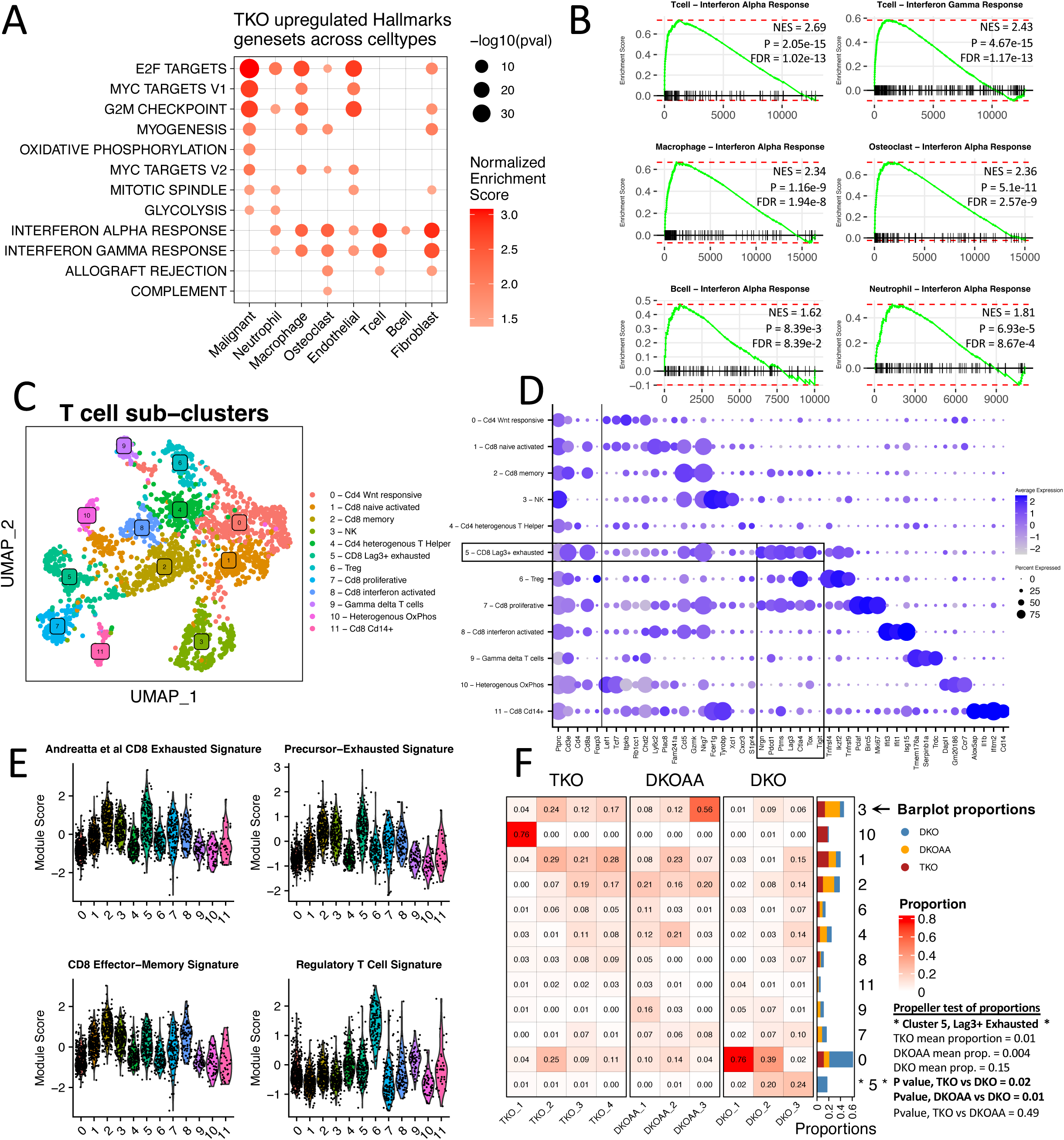
Induced immune activation in the form of interferon pathway activity and reduction of T cell exhaustion. A: Dotplot of Hallmarks gene sets significantly upregulated in TKO relative to DKO across cell types. B: GSEA plots showing enrichment of interferon response pathways in genes differentially expressed in TKO relative to DKO among immune cells. C: UMAP showing sub-clustering of T cells. D: Canonical and data-driven markers of T cell sub-clusters. E: Signature gene scores of T cell states derived from marker genes in a published meta-analysis of tumor-infiltrating T cells^34^. F: Compositional analysis of T cell sub-clusters across 3 OS tumors.

One potential mechanism underlying the increased immune activity in TKO tumors may be related to cellular stress response, as endoplasmic reticulum stress and replicative stress have both been linked to interferon signalling^31,32^. We therefore checked for differential expression of pathways related to stress and unfolded protein response [**Supplementary Figure 8**]. Surprisingly, we detected significant upregulation of unfolded protein response in malignant cells and endothelial cells in TKO and DKOAA. However, TKO macrophages appeared to downregulate this pathway while DKOAA macrophages showed upregulation of the pathway. These findings underscore the value of scRNA-seq analysis, uncovering cell type specific effects that could be missed in the analysis of bulk tumors. Interestingly, we also observed upregulation of gene sets related to replicative stress (including genes such as *Clspn* and *Chek1*) in malignant cells, macrophages, and endothelial cells, as well as telomere stress (with genes such as *Acd*) in malignant cells. These findings suggest a link between the hyper-replication phenotype observed in *Skp2* knockout and inflammation via induction of cellular stress in malignant cells and macrophages. Given these findings, further exploration of the mechanistic relationship between *SKP2*, endoplasmic reticulum stress, and replicative stress is warranted.

We next investigated genes related to interferon signaling [**Supplementary Figure 9**]. We found that the expression of interferon ligands was sparse and restricted to specific cell types, while upstream regulators *Tmem173 (*STING*), Cgas,* and *Tbk1*, as well as interferon receptors were more uniformly detected. Nevertheless, we did observe evidence of modest upregulation of upstream pathway genes *Tmem173* among B cells and *Cgas* among endothelial cells, receptor gene *Ifnlr1* among neutrophils, and Type 1 interferon ligand *Ifna4* among macrophages in TKO relative to DKO (P < 0.05, EdgeR). We also performed cell-cell communication analysis using CellChat with our scRNA-seq data^33^. Interferon signaling was not detected by CellChat, likely due to low levels of the ligand expression, but it did detect a multitude of significant signaling differences across the three models [**Supplementary Figure 10].** Considering the complexity in different cell types and how they can crosstalk, more research will be required to study the mechanism underlying interferon pathway induction in the OS tumors.

To investigate how each cell type was affected by *SKP2* disruption in a higher resolution, we carried out sub-clustering analysis using batch-corrected data (see **Methods**). Various transcriptomically distinct cellular sub-populations were discovered and their corresponding markers identified. [**Supplementary Figures 11 and 12, Supplementary Table 7]**. For tumor infiltrating T cells, subclustering distinguished 11 T cell subtypes and additionally a separate cluster expressing NK markers [**Fig 3C**]. As in the full dataset with all cells, we annotated these subclusters based on canonical markers in literature and data-derived top marker genes [**Fig 3D**]. Notably, this uncovered a CD8 T cell subcluster (cluster 5) characterized by high expression of dysfunctional T cell “exhaustion” markers, such as PD1 (*Pdcd1*), *Ctla4*, *Tigit,* and especially *Lag3*, along with exhaustion regulator *Tox*. To corroborate our results, we obtained gene signatures from a prior meta-analysis^34^ of scRNA-seq datasets of tumor-infiltrating T cells and used them to compute signature scores for different T cell states, including a CD8 exhaustion signature, a closely related “precursor-exhausted” signature, a CD8 effector-memory signature, and a regulatory T cell signature [**Fig 3E**]. Concordant with our marker-based annotation, the results showed that the T cell subcluster 5, relative to all other subclusters, displayed a stronger expression of the “CD8 exhausted” (P < 0.001, Wilcox test) and “precursor-exhausted” (P < 0.001, Wilcox test) signature genes. While these exhaustion signatures were also high in subcluster 2, the top signature for that subcluster was “CD8 effector memory” [P < 0.001].

After annotating these T cell subclusters, we compared their abundances between DKO, DKOAA, and TKO tumors [**Fig 3F**]. Interestingly, the only significant difference was a reduction in the proportion of subcluster 5, the CD8 *Lag3*+ exhausted T cells, in both TKO and DKOAA compared to DKO. While CD8 exhausted T cells accounted for a large proportion of tumor-infiltrating T cells in DKO tumors, they were almost completely absent from TKO and DKOAA tumors.

Several other cell types also exhibited significant subtype alterations among DKO, DKOAA and TKO [**Supplementary Figure 13**, **Supplementary Table 8]**. These included *Naalad*2+ malignant subcluster 4, significantly depleted from both TKO and DKOAA tumors relative to DKO; *Camp+*/*Fcnb-* neutrophil subcluster 2, enriched in DKOAA relative to DKO; *C1qa+* osteoclast subcluster 0, enriched in DKOAA relative to DKO; *Chil3+* osteoclast subcluster 4, depleted from DKOAA relative to DKO; *Kcne3*+ endothelial cluster 6, enriched in DKOAA relative to both TKO and DKO; *Hs3t1*+ B cell cluster 2, depleted in TKO relative to DKO; and *Ighd*+ B cell cluster 1, enriched in DKOAA relative to DKO. While these results suggest a strong cell population difference in the OS tumors with disrupted *Skp2* function, the functional importance and clinical relevance of our observations will require more validation and further study.

Taken together, our results indicate that immune activation in OS, in the form of induction of interferon activity and downregulation of T cell exhaustion, is likely a key consequence of *SKP2* function disruption. The mechanism underlying immune activation may be related to induction of cellular stress in TKO and DKOAA malignant or microenvironment cells. Additionally, the concordance of many of these results between the TKO and DKOAA OS tumors indicates that the mechanism underlying this immune activation involves the function of p27.

### E2f activity was upregulated in TKO and DKOAA malignant cells

*Skp2* knockout has been shown to effectively prevent tumorigenesis in other cancers driven by p53 and Rb1 inactivation such as prostate and pituitary cancers, via E2F1-driven apoptosis^13^. Consistent with these reports, E2f target upregulation was observed in TKO and DKOAA malignant cells [**Fig 2A, 4B**]. To further investigate this, we applied SCENIC to predict and compare the activity of the E2F family of transcription factors [**Fig 4A, Supplementary Figure 14**]. From our scRNA-seq data, we were able to infer SCENIC regulon activity scores for *E2f1*, *E2f2*, *E2f7*, and *E2f8*. Among malignant cells, the regulon scores of *E2f1* (P = 0.03), *E2f7* (P = 0.02), and *E2f8* (P = 0.03) were significantly greater in TKO than in DKO, while only *E2f8* was significantly higher in DKOAA than in DKO (P = 0.03, T-test). Nevertheless, numerous target genes of the E2F family showed upregulation in TKO and DKOAA relative to DKO malignant cells [**Supplementary Figure 14 G-J**]. Additionally, we investigated cell cycle phases and uncovered evidence of cell cycling among malignant cells and osteoclasts, but to a lesser degree in other cell types [**Supplementary Figure 14D**]. Comparison of S phase and G2/M phase score distributions revealed an increase in S phase among malignant cells compared to G2/M, while both were upregulated in TKO and DKOAA relative to DKO. The differences of these scores were less consistent in other cell types across the three OS models [**Supplementary Figure 14E-F**]. A critical link between *SKP2* disruption, E2f hyperactivation, and over-replication of DNA was reported to explain the induction of apoptosis upon *Skp2* KO^13^. Consistent with this, we found significant upregulation of an apoptosis gene set, including genes such as *Bbc3* and *Bid*, in both TKO and DKOAA relative to DKO malignant cells [**Fig 4B, Supplementary Figure 14L**].

**Figure 4:**
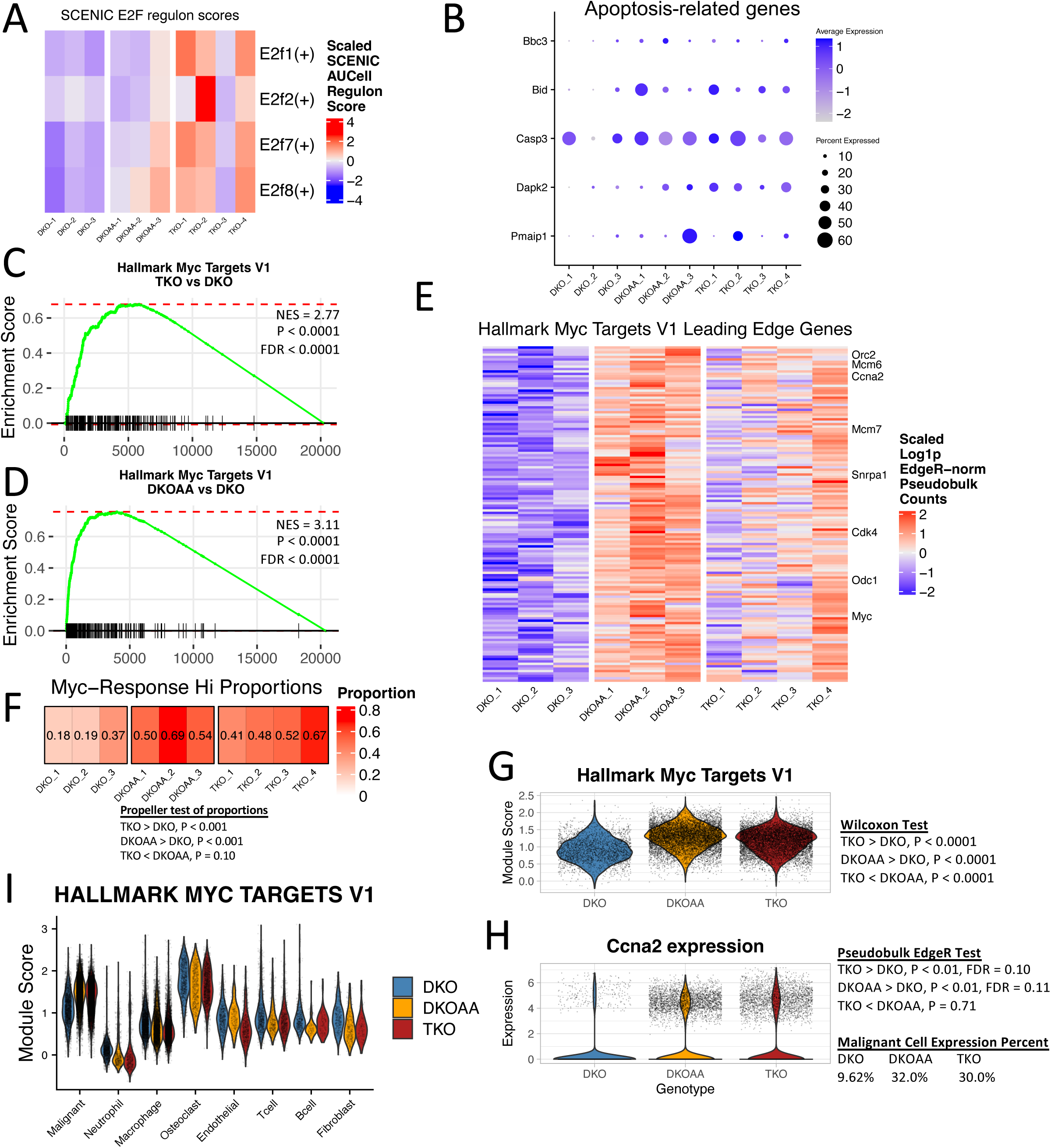
Upregulated Myc target activity in TKO and DKOAA malignant cells. A: SCENIC transcription factor activity scores of E2F family transcription factors. B: Expression of apoptosis-related genes derived from the Reactome Apoptosis gene set. C,D: GSEA plot of one of the “Hallmark Myc Targets V1” gene set in TKO and DKOAA relative to DKO, respectively. E: Heatmap of GSEA leading-edge genes in the “Hallmark Myc Targets V1” gene set. The intersect of leading-edge genes from the TKO and DKOAA enrichments is shown. F: Compositional analysis of *Myc*-high responder cells. G: Violin plot for malignant cells only of signature scores calculated from genes in the “Hallmark Myc Targets V1” gene set. H: Violin plot for malignant cells of *Ccna2*, a *Myc* target gene. I: Violin plot of the *Myc* signature scores in all cells.

Taken together, our scRNA-seq analysis supports the established relationship between *Skp2* functional loss, E2f induction, and apoptosis. The increase of apoptosis and the higher activity of immune microenvironment (described above) may contribute to slow tumor growth and small tumor size in TKO relative to DKO mice^17,18^.

### *Myc* activity was upregulated in TKO and DKOAA malignant cells

While activation of E2F targets seems advantageous by induction of tumor cell apoptosis, *Myc* targets, which are often considered to drive tumor development, were also significantly upregulated in both TKO and DKOAA malignant cells relative to DKO [**Fig 4C,D**]. These included metabolism-related genes such as *Odc1*, proliferation-related genes like *Ccna2* and *Cdk4,* and *Myc* (*c-Myc*) itself [**Fig 4E**]. To further investigate this, we used the “Hallmark Myc Targets V1” gene set to compute a signature score and defined cells above the median score as “High Myc-responder” cells. The proportion of such cells was significantly higher in TKO and DKOAA than in DKO [**Fig 4F**]. At the single cell level, the score distribution was greater in TKO and DKOAA than in DKO [**Fig 4G**]. More TKO and DKOAA cells exhibited strong expression of *Ccna2*, a canonical *Myc* target, than DKO cells as well. However, malignant cells, along with osteoclasts, expressed *Myc* targets at a level relatively higher than most non-malignant cell types, regardless of model (including DKO), suggesting that *Skp2* functional loss may amplify the *Myc* target transcriptional program that is already activated in the malignant lineage [**Fig 4I, Supplementary Figure 14M**]. We also studied *Myc* regulon activity using SCENIC but found only a modest upregulation. This may be due to some difference in the gene sets defined for *Myc* targets between the Hallmark and the SCENIC databases. However, examination of the expression patterns of the *Myc* target genes from SCENIC revealed a similar pattern as those observed in the “Hallmark Myc Targets V1” gene set [**Supplementary Figure 14C, K].** Alternatively, the up-regulation of *Myc* targets may be driven by other transcription factors.

Given the lack of impact of *Skp2* knockout in *Myc*-driven Burkitt lymphoma^19^, our findings suggest that the upregulation in TKO may be a mechanism adopted by OS tumor to escape from the lack of *Skp2.* Future studies will be required to address whether there is a direct link between *Skp2* disruption, *Myc* target induction and persistent (albeit delayed) tumorigenesis in TKO and DKOAA; but *Myc* targets nevertheless represents a potential target for synergistic therapy with *SKP2* inhibition.

### *Skp2* knockout places selective pressure on OS malignant cells

In addition to *Myc* target activation, we also wanted to explore additional mechanisms that may contribute to escape from *Skp2* functional disruption, leading to tumor formation in the TKO and DKOAA mice. OS is notorious for its complex genomic instability^35,36^, which may provide an evolutionary mechanism by which OS tumors resist treatment^37^. This may also be related to the frequent *RB1* and *TP53* loss in OS. To study if *Skp2* disruption could further increase genome instability in the OS, and thus allow tumor development, we inferred CNV profiles from scRNA-seq data and compared them across the three OS models. We found large variations among all the tumor samples [**Fig 5A**], but some consistent differences among malignant cells of the three OS models were observed. The number of genes affected by extreme amplifications (defined as 2X copies or more) was significantly greater in TKO and DKOAA compared to DKO malignant cells (P < 0.0001 in both, Wilcoxon test) [**Fig 5B, top right panel**]. However, the number of genes predicted to have homozygous deletions was significantly reduced in TKO and DKOAA compared to DKO (P < 0.0001 in both) [**Fig 5B, lower right panel**]. TKO malignant cells had more genes affected by both extreme amplifications and deletions relative to DKOAA (P < 0.0001 in both). To study this at single cell level, we performed PCA on the malignant cells, based on what genes were affected by the inferred CNVs. Interestingly, this revealed a greatly increased CNV heterogeneity in TKO and DKOAA than in DKO [**Fig 5C**]. Further study of the PCA embeddings demonstrated that DKO tumor samples appeared quite similar to one another, while TKO and especially DKOAA malignant samples had extremely diverse CNV profiles from DKO and from one another [**Fig 5D**].

**Fig 5:**
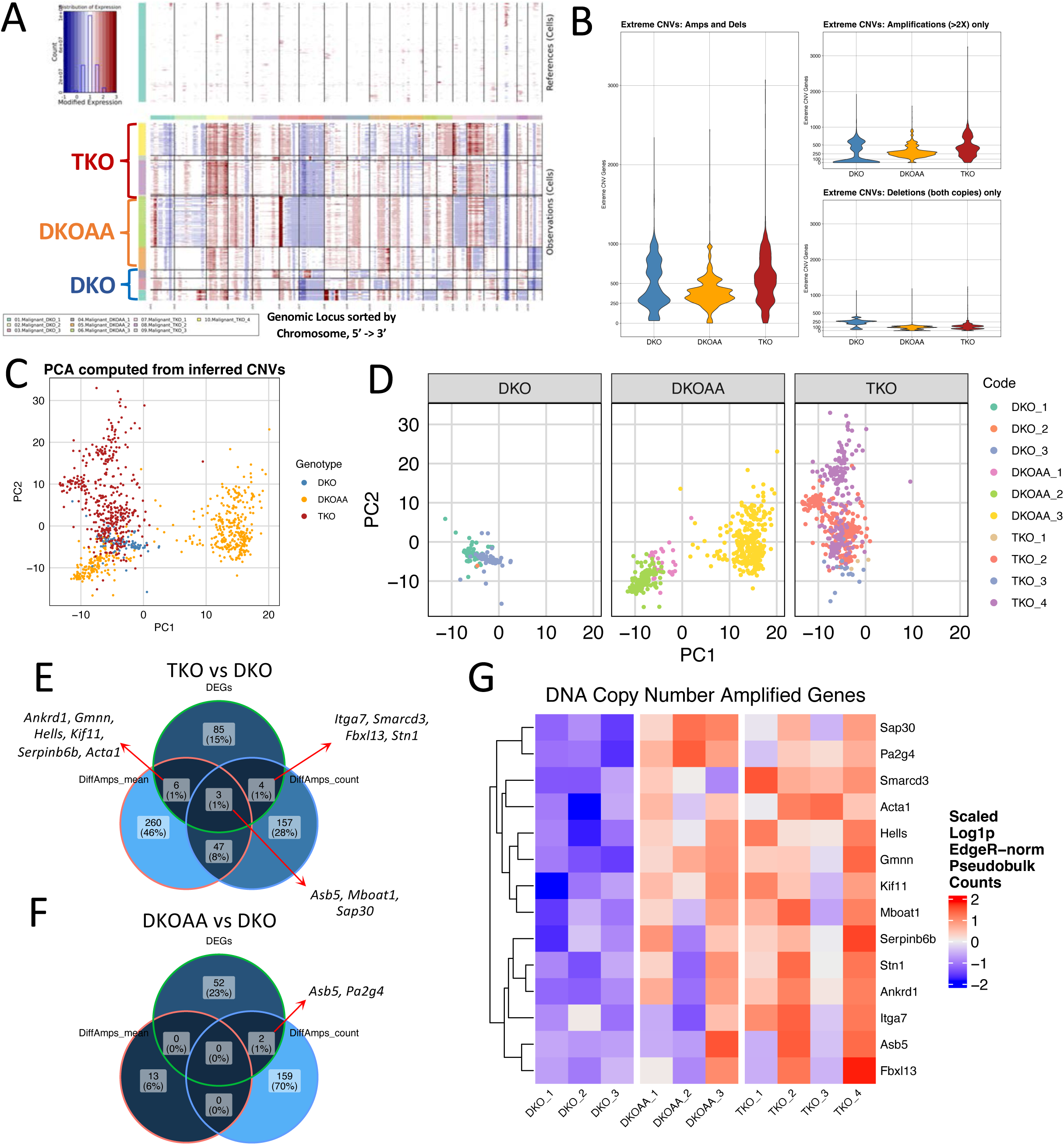
Genome instability in OS malignant cells. A: Heatmap showing Hidden Markov Model results of CNVs as determined by InferCNV. The top heatmap shows CNV states in macrophages (reference non-mutated cells), while the bottom heatmap shows malignant cells from each sample (rows). The columns represent chromosomes. B: Violin plots showing the number of genes affected by “extreme CNVs” (2x deletions or 2x> amplification). The left panel shows all extreme CNVs, the top-right panel shows extreme amplifications, and the bottom-right panel shows extreme deletions. C: PCA plot of malignant cells using inferred CNVs. D: Same PCA split but by OS models and colored by samples. E: Venn Diagram showing comparison of two differential CNV identification methods with differential expressed genes of TKO vs DKO. F: Venn Diagram showing comparison of two differential CNV identification methods with differential expressed genes of DKOAA vs DKO. G: Heatmap showing expression patterns of significantly DNA copy number amplified genes that were also significantly overexpressed in TKO and DKOAA.

We next investigated if specific genes affected by CNVs could be identified among DKO, DKOAA and TKO, despite high within-group variability. We devised two replicate-aware, pseudobulk-based methods for comparison of CNVs across the models, the first involving comparison of mean inferred CNV states, and the second counting up CNVs and comparing proportions [**see Methods**]. While we did not detect any significantly differentially deleted genes, we found some genes significantly more amplified in (multiple) TKO and DKOAA tumors relative to DKO, some of which were also significantly overexpressed in TKO and DKOAA malignant cells [**Fig 5E, 5F**]. More such genes were detected from TKO compared to DKOAA, which may be a result of the high variability observed among DKOAA samples.

Among the genes found to be both significantly amplified and overexpressed in TKO and DKOAA relative to DKO [**Fig 5G**], two genes were especially striking, *Fbxl13* and *Asb5*, two poorly studied E3 ligase substrate-targeting genes. *Fbxl13* is a member of the same Leucine-rich F-box family as *Skp2* (which is also known as *Fbxl1*) and also functions as a substrate-recognizing component of the SCF complex. It has been shown to target the centrosome protein *CEP192* for degradation, linking this gene to functions including cell motility and potentially mitosis^38^. *Asb5* is a poorly studied gene in the ankyrin repeat and SOCS-box containing protein family, many of which have been shown to cooperate with a different E3 ligase complex involving the scaffold protein *Cul5* via the SOCS domain (as opposed to *Cul1* in the SCF^SKP2^ complex)^39^. Other noticeable amplified and overexpressed genes in TKO include *Mboat1*, a negative regulator of ferroptotic cell death^40^; *Sap30*, a histone deacetylase associated with worse prognosis in neuroblastoma^41^; several genes related to regulation of DNA replication including *Stn1*, *Hells,* and *Geminin*; as well as *Itga7* and SWI/SWF (BAF complex) component *Smarcd3*, both of which have been linked to cancer stemness^42,43^. In DKOAA, besides *Asb5*, we observed amplification of the gene *Pa2g4*, which has been shown to cooperate with histone deacetylases to downregulate *E2f1* target activation^44^. Additionally, while the ankyrin-repeat containing transcription factor *Ankrd1* has previously been shown to be upregulated by *SKP2* in a manner dependent upon non-proteolytic ubiquitination of YAP and would thus be expected to be downregulated in TKO, we observed both overexpression and amplification of this gene in TKO, potentially indicating context-dependent mechanisms of regulation in OS^45^. The functional relevance of these genes, and whether there may be target redundancy between *Asb5*, *Fbxl13*, and *Skp2* warrants further studies.

These results indicate that the mutation spectrum of OS tumors is strongly perturbed by *Skp2* knockout in a manner particularly driven by copy number amplifications. While high variability existed among the malignant cell population, evidence for amplification in some cells was detected among several E3 ligase substrate-targeting genes with potentially similar function like *Skp2*, along with other genes related to cancer stemness, DNA replication, and cell death inhibition. These results suggest that *SKP2* functional disruption may place strong selective pressure or establish a genomic context for OS to evolve escape mechanisms.

### Mouse models of OS predict improved survival of *SKP2* disrupted interferon response activity

As mentioned above, the cell types present in our immunocompetent autochthonous murine OS tumors are similar to those reported from patient tumors [**Fig 1B**]^20,21^. To further investigate how well our OS models recapitulated human disease, we downloaded and integrated data from 17 human patient tumors [**Fig 6A**] ^20,21^, which included 11 pre-treated, resected samples (“NatComm_2020”) and 6 treatment-naïve, upfront-resected samples (“FrontOncol_2021”). Cell-level annotations were not provided by either study, so we re-annotated cell types after performing cell clustering, marker gene identification, and InferCNV analysis of the integrated data [**Fig 6B, Supplementary Figure 15, Supplementary Table 9]**. As in murine tumors, malignant cells from patient tumors have a much higher number of genes affected by 2X deletions or amplifications than stromal or immune cells [**Fig 6C**]. Additionally, cell type marker genes in human tumors showed expression patterns very similar to those in mouse tumors [**Fig 6D**]. To better quantify the transcriptomic similarity of cell types between human and mouse OS, we identified top marker genes for individual cell types in the mouse OS and used them to compute “cell type scores” in the human data. The results indicated high specificity of murine-derived cell type gene signatures [**Fig 6E**].

**Figure 6:**
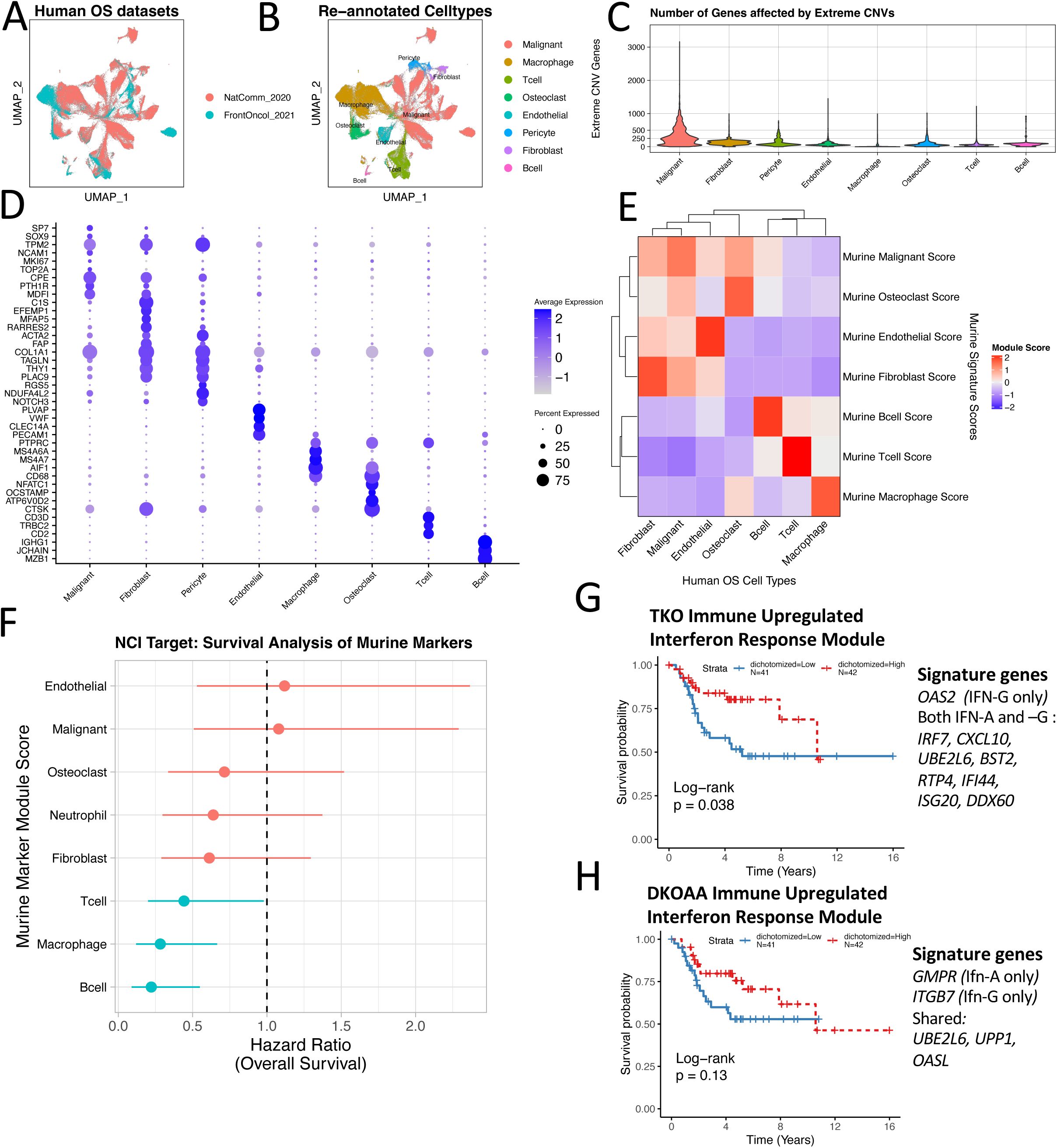
Relating mouse OS models to patients. A,B: UMAPs of human OS patient scRNA-seq data integrated from two publications, colored by data sources (A) and cell types (B). C. Violin plot showing the number of genes affected by extreme CNVs (>= 2x deletion or amplification) across cell types. D: Canonical and data-driven markers of cell types in integrated human OS. E: Heatmap showing expression scores of murine OS celltype markers in patient tumor cells. Rows correspond to murine marker genes, and columns refer to human OS cell types. F: Forest plot illustrating survival analysis of murine cell type signatures in the NCI Target OS cohort. G: Kaplan-Meier (KM) plot showing survival association of an expression score calculated from genes in the Hallmark Interferon-Alpha and Interferon-Gamma Response gene sets that were also upregulated in TKO immune cells relative to DKO. H: KM plot showing survival association of an expression score calculated from genes in the Hallmark Interferon-Alpha and Interferon-Gamma Response gene sets that were also upregulated in DKOAA immune cells relative to DKO.

Some cell types, however, were detected in only human or mouse OS, including pericytes in human tumors, and neutrophils in mouse tumors. Technical factors in sample preparations likely explain this difference. Pericytes may have been insufficiently sampled in murine tumors due to lower sample size (10 mouse tumors versus 17 human tumors), while neutrophils are known to be difficult to capture via scRNA-seq^46^ and laboratory conditions may be more suitable for such challenging cells compared to the operating room.

Considering this transcriptomic similarity between murine and patient tumors, we decided to conduct survival analysis in a larger cohort of patients with bulk RNA-seq data, the NCI-TARGET cohort, to test the impact of cell types. We observed significant positive survival benefits in the patient samples with greater expression scores for the markers of murine T cells, macrophages, and B cells [**Fig 6F**], suggesting increased abundance of these immune cell types is beneficial to patients. We next studied if any pathways significantly differentially expressed upon *Skp2* functional disruption were associated with survival. Starting from the Hallmarks Interferon (Alpha- and Gamma-) Response gene sets, we selected the genes upregulated in TKO and DKOAA immune cells to calculate gene scores for the samples in the NCI TARGET cohort. We found that high expression of both were associated with improved survival [**Fig 6G**, **Fig 6H**].

Taken together, these results indicate that the tumor microenvironment and transcriptome of our murine models of OS resemble those of patient tumors, and that immune infiltration is associated with improved survival in human OS. Furthermore, the interferon response gene signature up-regulated in *SKP2*-disrupted OS is correlated with improved survival in OS patients.

### Pathologic subtype and lineage infidelity influences OS tumor heterogeneity

Patient OS can present in several pathologic subtypes including osteoblastic OS, chondroblastic OS characterized by increased chondroid matrix, and fibroblastic OS characterized by lower levels of matrix production^47^. The first scRNA-seq study of human OS reported some clear differences between osteoblastic and chondroblastic OS, with the latter characterized by expression of canonical chondroblast markers including *COL2A1*, *ACAN,* and lineage-driver *Sox9,* reduced presence of osteoclasts in the tumor microenvironment, as well as active differentiation to osteoblastic cell state^20^. We wondered if some levels of tumor heterogeneity in the murine data [**Fig 7A**] could be related to pathologic subtypes although we did not expect *Skp2* knockout would shift OS subtypes.

**Fig 7:**
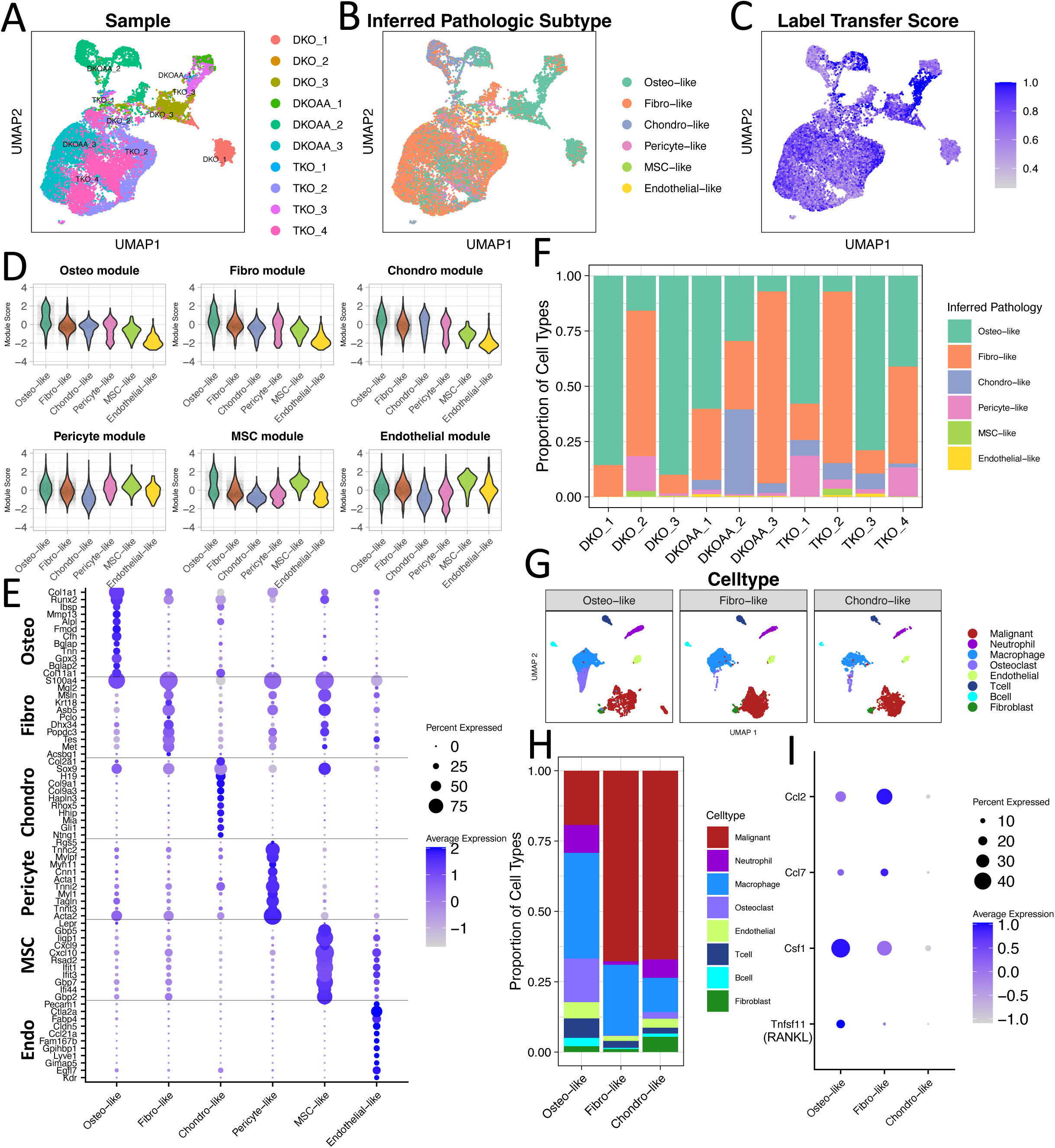
Pathologic subtype and lineage infidelity in mouse OS tumor. A,B: UMAP showing malignant cells from OS tumors, colored by sample (A) and pathologic subtype (B) as inferred via label transfer from a murine non-malignant bone atlas dataset. C: The same UMAP, colored by label transfer scores. D: Violin plots of signature scores computed for markers of various bone cells. E: Dot plot of markers of malignant subtype classifications. F: Bar plot showing proportions of malignant cells annotated to each pathologic subtype in each sample. The subtype with the highest proportion is considered the inferred pathologic classification of the indicated tumor. G: UMAP of all OS tumor cells similar to Fig 1B but split according to sample-wise inferred pathologic classification. H: Proportion of cell types in each pathologic subtype. I. Expression patterns of genes related to macrophage and osteoclast differentiation among malignant cells from each subtype.

To study this, we relied on a published scRNA-seq atlas of non-malignant bone and bone marrow niche cells^48^, because it contained various clusters of osteo-lineage cells (osteoblasts), fibroblasts, chondroblasts, along with pericytes, endothelial cells, and “mesenchymal stromal cells” (MSCs), which express hematopoietic niche factors such as *CXCL12* and stem cell marker *LEPR*. For the purpose of our analysis, we merged sub-clusters of the same cell types to obtain clusters related to broad cell types [**Supplementary Figure 16**].

We performed a label transfer using Seurat, in which malignant OS and the non-malignant atlas bone cells were integrated and cell labels were estimated based on transcriptomic similarity. The results indicated that OS malignant cells were composed mostly of osteo-like, fibro-like, and chondro-like cells [**Fig 7B**]. Additionally, there were small numbers of cells annotated as pericyte-like, MSC-like, and endothelial-like. Inspection of label transfer score, a measure of transcriptomic similarity between malignant and matching bone atlas cells, revealed high score appeared to correlate with cluster areas with high cell type purity [**Fig 7C**].

We also applied label transfer using the same bone atlas to our non-malignant endothelial and cancer-associated fibroblast cell types [**Supplementary Figure 17**]. These revealed an almost perfect match for endothelial cells and a high score for most fibroblast cells. Interestingly, some of the fibro-cells were called as osteo-like, which likely explained the lower label transfer scores. This raises the possibility that non-malignant osteoblast cells may be present in the OS tumor microenvironment, alongside (and transcriptomically similar to) cancer-associated fibroblasts.

To further establish the pathologic classifications, we derived markers from the simplified bone atlas dataset for computing signature scores in the malignant OS cells [**Fig 7D**]. This revealed that while osteo-like and chondro-like cells strongly express the osteo and chondro signatures, respectively, the fibro-like cells had only a modest upregulation of the fibro signature. The minor cell types including pericyte-like, MSC-like, and endothelial-like OS cells all had robust upregulation of their respective signature genes. Notably, the osteo-like cells appeared to also express every cell type signature, indicating perhaps more plasticity.

We also inspected the malignant OS subtype markers, along with some canonical cell type markers, and observed fairly high expression specificity for most cell types, with the exception of the fibro-like markers, which were expressed somewhat more promiscuously [**Fig 7E**]. Restriction of marker analysis to the three main subtypes (osteo-like, fibro-like, and chondro-like) revealed more specific markers [**Supplementary Figure 18A**]. Using SCENIC, we also uncovered numerous transcription factors specific to the main pathologic subtypes [**Supplementary Figure 18B**]. We noticed that *Asb5* was also found to be overexpressed in the fibroblastic subtype. In concordance with other results, *Asb5* was found to be almost completely silent in non-malignant subtypes, but was upregulated in TKO and DKOAA in all malignant subtypes [**Supplementary Figure 18C,D]**.

Using the inferred pathologic subtypes, we called each tumor sample as either osteo-like, fibro-like, or chondro-like based on which subtypes had the highest frequency in the malignant cells [**Fig 7F**]. Osteo-like samples included DKO_1 (86% osteo-like cells), DKO_3 (90%), DKOAA_1 (60%), TKO_1 (58%), and TKO_3 (79%); only one sample was classified as chondro-like, DKOAA_2 (38%); and fibro-like samples included DKO_2 (66%), DKOAA_3 (87%), TKO_2 (77%), and TKO_4 (44%). DKOAA_2, the only sample classified as chondro-like, was in fact highly mixed and contained high proportions of all three types (38% chondro-like, 30% osteo-like, 31% fibro-like, and small proportions of others). While at first we considered this sample as too highly mixed to classify, we decided to keep this result, because the data matched the characteristics of chondroblastic OS patient tumors, where sub-clusters of intermediate cells transdifferentiating between chondro- and osteo-like phenotypes were found ^20^.

The OS subtype analysis is important because it uncovered a striking difference in the makeup of the tumor microenvironment [**Fig 7G**]. Comparison of the cell type proportions in the microenvironment showed a clear depletion of osteoclast cells in fibroblastic tumors [**Fig 7H**]. Osteoclasts, macrophage-like hematopoietic cells involved in bone resorption, contributed a large proportion of cells in the osteo-like tumors (16%) but represented a minor proportion in chondro-like tumors (2%) and were exceedingly rare in fibro-like tumors (<0.1%) (Osteo vs fibro P = 0.003, propeller test; chondro had just one sample). We noted that osteoclasts exhibited cycling characteristics [**Supplementary Figure 14D**], indicative of active osteoclastogenesis in the tumor microenvironment, as in human tumors^20,49^. Based on this, we inspected cytokines involved in this process and observed decreased expression of the macrophage maturation cytokine *Csf1* and osteoclast differentiation factor RANKL (*Tnfsf11*) in the fibro-like and chondro-like malignant cells relative to the osteo-like cells, while monocyte homing cytokines such as *Ccl2* and *Ccl7* were not changed [**Fig 7I**]. This indicates that the rarity of osteoclasts in the chondro- and especially fibro-like tumors may be a result of reduced tumor-mediated osteoclastogenesis, rather than reduced general monocyte infiltration. Although this demonstrates a clear cell compositional difference among the tumor microenvironment between the OS subtypes, the small sample size prevents us from making any clinical or functional implications, but it could be important for future study.

Given these findings, we performed differential expression analysis across OS models stratified by pathologic subtype where samples were available, osteo-like and fibro-like. Due to low sample sizes, we were forced to rely on non-pseudobulk methods which may be susceptible to false positives; we applied the Wilcox test. Overall, we found similarity with the non-stratified analysis, including upregulation of *Myc* targets among malignant cells in TKO and DKOAA versus DKO in both osteo- and fibro-like tumors [**Supplementary Figure 19**]. However, one notable exception was that EMT-related gene expression was found to be upregulated in TKO and DKOAA osteo-like tumors. Inspection of EMT-related markers indicated that this seemed to be driven particularly by increased expression in TKO of ECM-related genes such as *Mmp23, Pcolce, Timp1, Sparc,* and *Col1a1*, while most genes associated with motility remained downregulated in TKO [**Supplementary Figure 20**].

Taken together, these results indicate that Osx-cre driven transgenic OS can develop to distinct OS subtypes as in human OS. Our analysis also provides transcriptomic markers of these subtypes that can be evaluated in patient OS. Furthermore, the OS subtype appears to influence the microenvironment cell composition, e.g., depletion of osteoclasts from chondroblastic tumors as reported in patients, and even more pronounced in fibroblastic OS. Validations of the inferred subtypes and the corresponding microenvironment difference are needed to confirm our findings, however, our findings indicate that pathologic subtype is a strong source of molecular variation in OS tumors that should be considered during sampling in future studies.

### Upregulation of a hazardous myogenic transcriptional program in OS with *Skp2* disruption

In general, we observed high concordance in the results of comparing TKO and DKOAA to DKO. This indicates that a p27-dependent mechanism is critical for *Skp2* function. However, there are differences between TKO and DKOAA models. For example, TKO malignant cells upregulated genes related to myogenesis relative to DKO, while DKOAA did not [**Fig 2A, 2B**]. We thus made a direct comparison between TKO and DKOAA and confirmed upregulation of myogenesis genes in TKO relative to DKOAA. [**Fig 8A,B**]. Genes driving this enrichment included transcription factors (e.g., *Mef2a*) and muscle function genes like *Myl1* and *Tnnc2*, components of the sarcomere, an organelle in striated muscle [**Fig 8C**].

**Figure 8:**
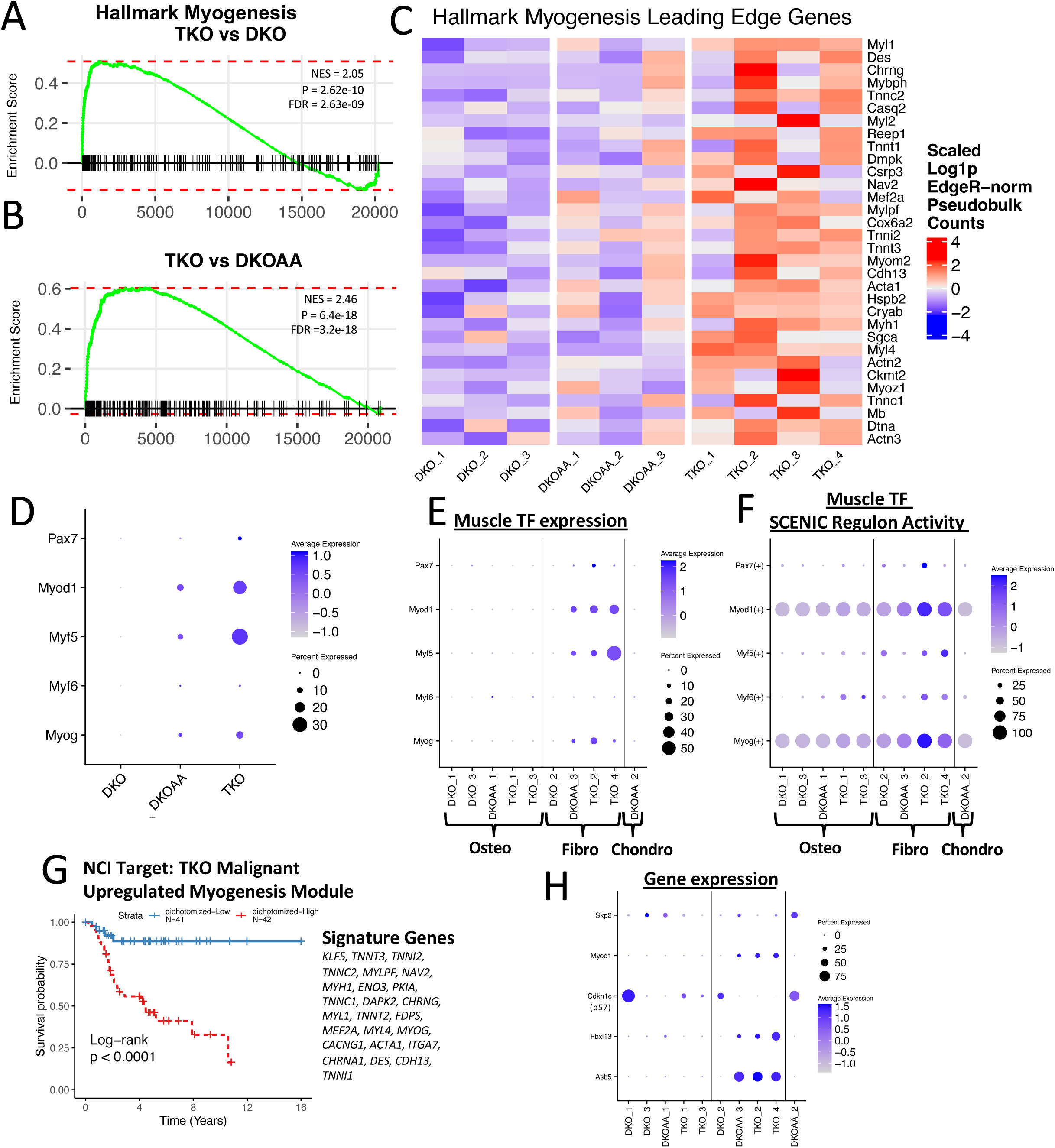
Upregulation of a hazardous myogenic transcriptional program in TKO. A-B: GSEA plots showing enrichment of the Hallmark Myogenesis gene set in TKO vs DKO and TKO vs DKOAA respectively. C: Heatmap showing leading edge genes from the Hallmark Myogenesis gene set from GSEA enrichment of TKO vs DKO. D: Dot plot showing expression of canonical muscle transcription factors. E: Dot plot showing expression of muscle transcription factors by samples, sorted by inferred pathologic subtypes. F: Dot plots showing “regulon activity” as inferred by SCENIC. G: KM plot showing survival association of an expression score calculated from genes in the Hallmark Myogenesis gene set that were also upregulated in TKO malignant cells relative to DKO. H: Dot plot showing expression profiles of key genes related to myogenic signature.

To confirm this, we examined the expression of transcription factors key for muscle differentiation, myogenic master regulator *Pax7* and the four myogenic regulatory factors *Myod1*, *Myf5, Myf6*, and *Myog*. We found that they were also upregulated in TKO [**Fig 8E**]. SCENIC analysis further confirmed that targets of these transcription factors were more active in TKO [**Fig 8F**]. Interestingly, there seemed to be a pattern of upregulation in some inferred pathologic subtypes. Fibro-like samples TKO_2 and TKO_4 had the strongest expression and SCENIC regulon scores, followed by DKOAA_2.

We were intrigued by this finding and decided to investigate the clinical relevance of this ectopic myogenesis program. Analyzing the NCI Target OS cohort, we observed a dismal prognosis in patients highly expressing the TKO myogenesis gene set [**Fig 8G**], when the TKO malignant upregulated genes that were also present in the Hallmark Myogenesis gene set were used to compute myogenesis scores for stratifying the patient cohort. In the OS patient scRNA-seq data, most of these genes were specifically expressed by malignant cells, but were restricted to one cluster, indicating the program involves a strongly distinct transcriptomic phenotype in OS [**Supplementary Figure 21**]. The human myogenic cluster (Malignant C23) was mostly composed of cells from one sample, “NatComm_2022 BC17”, a lung metastatic tumor. Additionally, the expression level of *SKP2* did not appear significantly lower than in other clusters, suggesting OS is capable of myogenic lineage plasticity in a manner that does not involve *SKP2*.

It has recently been reported that *SKP2* was upregulated by *MYOD1* in the context of rhabdomyosarcoma to drive a stem-like state of incomplete, non-differentiated myogenesis, via degradation of p57, thus linking *Skp2* with myogenesis in sarcoma^50^. We did not observe a clear pattern of transcriptional upregulation of p57 (*Cdkn1c*) in TKO or fibroblastic samples, but interestingly, we did detect upregulation of *Asb5* and *Fbxl13* in fibroblastic TKO and DKOAA samples [**Fig 8H**]. In the human OS scRNA-seq data, *ASB5* was also specific to the same malignant cluster that expressed the myogenic signature [**Supplementary Figure 21**]. *Asb5* expression has been detected in muscle stem cells before^51^. It is possible that p57 protein levels may be altered but not reflected at the transcript level and thus still involved in in the myogenic phenotype.

Overall, these results suggest that *Skp2* knockout unexpectedly led to upregulation of an abnormal myogenic pathway, perhaps a result of the malignant cells actively exploring evolution landscape. Since this program was also detected in the human myogenic OS cells expressing *SKP2*, it may not be directly regulated by SKP2.

## DISCUSSION

Here, we report the first scRNA-seq study of transgenic murine OS, as well as the first report of scRNA-seq in the context of *Skp2* knockout. Our findings indicate that the patient OS microenvironment is recapitulated by conditional ablation of *Rb1* and *Trp53* in the Osx lineage in mice. Our analysis also uncovered the two sides of targeting *SKP2* functions in OS. *Skp2* knockout or genetic disruption of its interaction with p27 resulted in increased immune activation, pro-apoptotic E2f activity, and reduced metastasis-related pathway activities. Using histologic examination of lungs, we validated the latter finding and found a strong reduction in OS metastasis after *Skp2* knockout. Conversely, our comprehensive analysis of the scRNA-seq data also implicates several potential mechanisms of resistance and escape from the loss of *Skp2*, which may explain why tumorigenesis is not entirely blocked but delayed in the TKO mice. Furthermore, we showed that *Skp2* knockout led to increased genome instability in the malignant OS cells without *Rb1*/*Trp53*, which may broaden the evolutionary landscape for OS to explore. Taken together, our study points to strategies that may be exploited for future development of synergistic *SKP2* targeted therapies.

We recently established a link between *Skp2* and immunomodulation and reported increased immune infiltration and interferon signaling in *Skp2* knockout tumors^18^. However, those results were obtained from analysis of bulk tumor tissues and thus could not resolve specific cell type effects. Here, we provide new evidence of widespread upregulation of interferon-response signaling in multiple cell populations within the OS microenvironment upon functional disruption of *Skp2*. Coupled with this, we uncovered evidence of reduced CD8 T cell exhaustion. Furthermore, we found evidence for increased replication stress and ER stress in several cell types after *Skp2* knockout, including malignant cells, macrophages, and endothelial cells. Interferon ligand expression was observed to be sparse, but we did detect upregulation of interferon ligands among macrophages. In addition to this, we observed upregulation of activity of the pleiotropic E2f family, including markers of increased cell entry to S phase along with malignant cell apoptosis. Further still, we observed evidence that the number and complexity of DNA copy number amplifications was increased in both TKO and DKOAA tumors. Taken together, our data supports a model in which *Skp2* knockout results in *E2f*-mediated overactivation of proliferative pathways related to DNA replication, resulting in cellular stress, increased mutation burden and possible neo-antigen formation, and induction of interferon activity [**Figure 9**]. The link between ER stress or replication stress with interferon has increasingly received strong interest as a novel target pathway for immunotherapy in several contexts^31,32^. Our findings support the notion that induction of cellular stress is a viable immunogenic target in OS and that *Skp2* disruption induces cell stress in a manner that may be exploited for cytotoxic immune activation.

**Figure 9:**
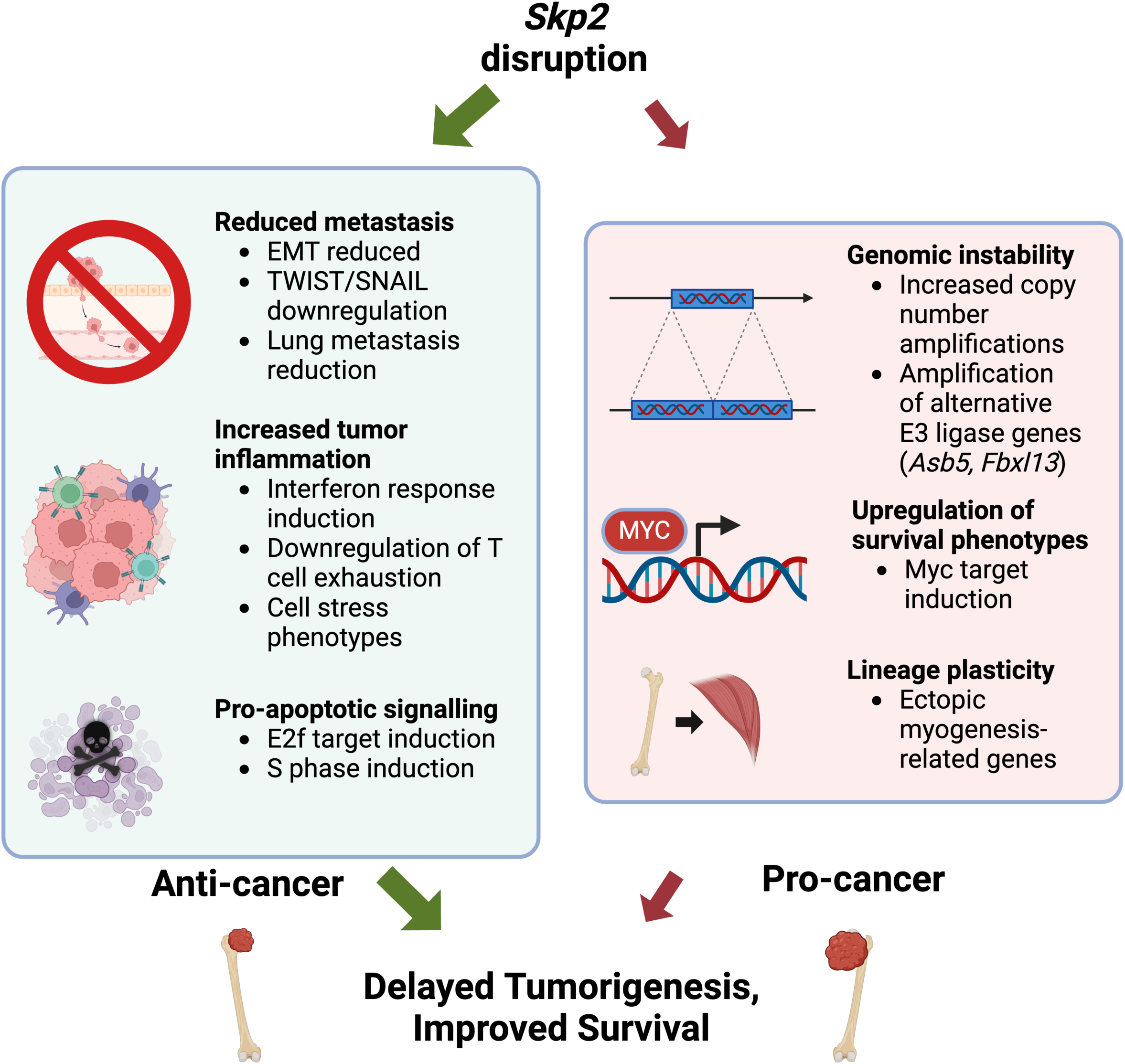
Working model summarizing main findings from mouse OS models. *Skp2* functional disruption led to differential expression of both anti-cancer and pro-cancer pathways, representing a battle between different forces in malignant cells and TMEs, which eventually result in delayed but not complete blockage of tumor development. Even the tumors escaped from *Skp2* disruption may experience the anti-cancer benefit of *Skp2* reduction, leading to slow and less aggressive tumors. Figure was created with BioRender.

In addition to these, we observed downregulation of metastasis-related signatures in TKO and DKOAA tumors, including downregulation of genes involved in epithelial-mesenchymal transition. This appears to be mainly driven by downregulation of genes related to extracellular matrix remodeling, cell motility, and associated transcription factors, rather than a shift in “epithelial” or “mesenchymal” cell lineage identity. Consistent with this, we observed a strong decrease in lung metastasis in TKO, but only a modest reduction in DKOAA mice. The divergent results between TKO and DKOAA indicates that the mechanism only partially involves the functional interaction between *Skp2* and p27. Alternatively, it is possible that the magnitude of p27 stabilization may differ between TKO and DKOAA, and the differences between models is thus p27 dosage-dependent. Nevertheless, taken together, this data supports the use of *Skp2* inhibition for suppressing lung metastasis in OS. One interesting future area of investigation would be to study if *Skp2* knockout reduces malignant “re-dissemination” from established metastatic tumors to novel sites (“metastasis to metastasis” spread). This has received increased interest as a mediator of disease burden in other tumor contexts such as breast cancer^52,53^.

Nevertheless, despite the strong survival benefits and delayed tumorigenesis after *Skp2* disruption, OS tumors from the TKO and DKOAA mice do eventually grow and progress. Our study points to several potential mechanisms that OS tumors may employ to escape the functional loss of *Skp2* [**Figure 9**]. Genetically, we detected strong mutational perturbation in TKO and DKOAA relative to DKO tumors, indicating that selective pressure took place, reminiscent of tumor evolution in the context of chemotherapy^37^. By analyzing CNV profiles inferred from the scRNA-seq data, we found that numerous genes gained copy numbers in TKO malignant cells, several of which also showed overexpression in TKO and DKOAA, including anti-apoptotic genes and understudied E3 ligase genes *Asb5* and *Fbxl13*. Further study, including whether these E3 ligases target substrates similar to those of *Skp2*, or whether these are targetable oncogenes in OS or other cancers, is warranted.

Additionally, we uncovered evidence of lineage plasticity among OS tumors by upregulation of an abnormal myogenic state. Although we did not have direct evidence for its contribution to OS development, interestingly high activation of the myogenic program was associated with poor survival in OS patients. Additionally, lineage plasticity is a mechanism commonly employed by cancer to escape treatment^54^. Notably, *ASB5* appeared to be correlated with OS muscle-related gene expression in both murine and human OS and has previously been observed to be expressed in muscle stem cells^51^. To our knowledge, this is the first report of malignant myogenic lineage plasticity in OS. The mechanism underlying this “osteo-myogenic transition” phenotype, including whether it may be driven by *ASB5*, or whether *ASB5* inhibition may reverse this state, remains an interesting open question. Furthermore, our previous analysis showed adipogenesis also appears to be affected by *Skp2* disruption^16,18^.

In addition to these possibilities, upregulation of *Myc* target activity in TKO and DKOAA tumors relative to DKO appears to be a strong candidate for escape from the loss of *Skp2*. This is further supported by a previous study reporting that *Skp2* knockout caused only a modest, non-significant improvement in survival in the context of murine models of *Myc*-driven Burkitt lymphoma^19^. Our data showed *Myc* upregulation in OS tumors but we did not detect amplification of *Myc* or any clear *Myc* targets. In addition, all malignant OS cells (including those from DKO) displayed higher *Myc* activity scores, indicating that targeting the *Myc* pathway is likely a good strategy for OS treatment and furthermore may increase the efficacy of *SKP2* inhibition. On the other hand, precision medicine approaches including genomics-informed application of CDK inhibitors have been proposed in OS and were efficacious in *MYC*-amplified patient-derived xenografts but did not completely stop tumor growth^55^. Future study of joint *SKP2* and *MYC* inhibition may thus reveal potent synergistic avenues of therapy.

OS is notorious for its extensive heterogeneity across patients. Our study demonstrated that mouse OS tumors were also characterized by strong inter-tumor molecular heterogeneity, as in patients. This heterogeneity may be driven by distinct mutational and CNV profiles as in patients, but we also found transcriptomic and microenvironment differences related to pathologic subtype. It has previously been shown that osteoclastogenesis appears to occur directly in the OS tumor microenvironment, and that osteoclasts may be rare in the microenvironment of chondroblastic OS^20^. Consistent with this, we observed evidence for osteoclastogenesis in murine OS and reduction of osteoclasts in chondroblastic tumors, but we also found almost complete ablation of osteoclasts from the microenvironment of fibroblastic OS, which was correlated with and may be mediated by reduced RANKL signaling. While the clinical implications of this remain unclear, this data suggests that the variability between OS histologic subtypes are strong enough that they should be considered during sampling and statistical power consideration in molecular studies. We also note that there appears to be an overrepresentation of the fibroblastic OS subtype among our samples, relative to its rare incidence in humans. It is possible that these tumors may be composed of softer cellular tissue compared to bony matrix found in osteoblastic tumors. The latter require lengthy collagenase treatment, which may reduce cell viability and be considered non-suitable for sequencing. If that is the case, our finding would represent a sampling bias toward the rare fibroblastic subtype. Alternatively, fibroblastic OS may be more common among this model of OS than in the patient population. Currently, our sample size is too small to address whether murine models of OS driven by Osx-cre ablation of *Trp53* and *Rb1* present with OS subtypes at similar proportions to those in human OS, or if the proportions are affected by *Skp2* disruption. Future studies could investigate whether alternative methods like single-nuclei RNA-sequencing may achieve better capture of cells from heavily mineralized osteoblastic OS tumors.

A key limitation of our study is the cross-sectional nature of the data, which prevents us from making any inferences about the temporal order of the various events found to be more or less active in the TKO and DKOAA tumors compared to the DKO. Expanding our analysis to samples collected from different OS growth stages would allow us to address whether the anti-cancer pathways (e.g., induction of immune activation) are active predominantly at the early stage while the pro-cancer pathways (e.g., *Myc* target upregulation) are induced at the later stage or concurrently [**Figure 9**]. Knowledge about the order is especially important for targeting SKP2 in OS patients. We should point out that TKO and DKOAA tumors in our study had already successfully developed strategies to cope with *Skp2* disruption partially. Another limitation is that our study was based on transcriptomics, and thus may miss post-transcriptional regulatory events. This is important because SKP2 regulates its targets at the protein level and many of the *Myc* and E2F targets are also regulated by protein abundance. Proteomic methods like mass spectrometry or CITE-seq could be used to address this in future studies. Additionally, we should mention that in patients OS tumors are removed and thus may not have the same resistance potentials as discussed for the mouse model.

In conclusion, our study supports the use of *SKP2* inhibition in OS treatment, but also reveals a complex battery of mechanisms that tumors may exploit for resistance. These suggest that a combination therapeutic strategy using *SKP2* inhibition and other drugs targeting the potential resistance pathways may be promising for OS treatment. In clinical practice, it will be important to know which of the potential resistance mechanisms are activated in individual patients. Our study provides biomarkers for molecular mechanisms associated with both effective anti-tumorigenic *Skp2* inhibition as well as oncogenic resistance pathways.

## MATERIALS and METHODS

### Establishment of animal models

Osx-Cre mice, *Rb1*^lox/lox^ mice, *Trp53*^lox/lox^ mice, *Skp2*^-/-^, and *p27*^T187A/T187A^ mice were described previously^15,16,56,57^. All mice used for experiments were on FVB, C57BL6J, and 129Sv hybrid backgrounds. First, Rb1^lox/lox^ mice were crossed with Trp53^lox/lox^ mice to generate Trp53^lox/lox^, Rb1^lox/lox^ mice, which were further crossed with Osx-Cre mice to generate Osx-Cre; Trp53^lox/lox^, Rb1^lox/lox^ mice (DKO). The Skp2^-/-^ mice were crossed with Osx-Cre;Trp53^lox/lox^,Rb1^lox/lox^mice to generate Osx-Cre;Rb1^lox/lox^;Trp53^lox/lox^; Skp2^-/-^ mice (TKO). The *p27*^T187A/T187A^ mice were crossed with Osx-Cre;Trp53^lox/lox^,Rb1^lox/lox^ mice to generate Osx-Cre;Rb1^lox/lox^;Trp53^lox/lox^; *p27*^T187A/T187A^ mice (DKOAA). Animals were maintained under a pathogen-free condition in the Albert Einstein College of Medicine animal facility, following animal experimental protocols reviewed and approved by Einstein Animal Care and Use Committee (#20180401), conforming to accepted standards of humane animal care. The tumor diameter was measured using a caliper every three days, and the relative tumor volume was calculated by the following formula: (length x width^2^) x 0.526. Tumors were resected when their volumes reached approximately 500 mm^3^.

### scRNA-seq data generation

Mice of each model presenting tumors close to 1.5cm in diameter were humanely euthanized. One tumor each from 4 TKO, 3 DKOAA, and 3 DKO animals was carefully dissected, removing surrounding tissues, and then rinsed thoroughly with cold PBS three times to eliminate impurities. Following the standard 10X Genomics sample preparation protocol, the samples were sectioned into 1–2mm pieces and subjected to digestion with type II collagenase and trypsin. To ensure complete digestion, the samples were incubated in a shaker at 37°C and shaken every 10 minutes. After 1.5 hours of digestion and incubation, the cell suspension was filtered through a strainer and centrifuged to remove the enzymes. Subsequently, red blood cell lysis buffer was added to the cell suspension to eliminate red blood cells. The number of cells was quantified using a Cell Counter (Bio-Rad, #TC-20), and only samples with an appropriate cell density (1,000 cells per μL) and a live-cell percentage greater than 85% were used for further single-cell RNA sequencing.

Cells were then loaded into a 10X Chromium instrument (10X Genomics) using the Chromium NextGEM Single Cell 3’ GEM, Library, and Gel Bead kit v3 at the Einstein Genomics Core. Sequencing of the DNA libraries was performed using an Illumina HiSeq 4000 system (Novogene, Sacramento, CA, USA) with 150 bp read length.

### scRNA-seq data analysis

#### Pre-processing and alignment

Sequencing reads in FASTQ format were demultiplexed and processed using CellRanger (v6.0.1; 10X Genomics) and aligned to the mm10 *Mus musculus* reference genome (mm10-2020-A; pre-prepared by 10X Genomics).

#### Filtering of RBCs, low-quality cells and doublets

We devised several methods for filtering of poor quality cells, which are described in detail at https://github.com/FerrenaAlexander/FerrenaSCRNAseq. In brief, we applied all filtering steps to each sample individually. For filtering of red blood cells and other low complexity cells, defined as cells with very few unique genes detected given the number of UMIs, we devised a two-step regression approach, in which a linear regression and LOESS regression model were fit with log(number of UMIs) as the predictor and log(number of unique genes) as the dependent variable. Barcodes with LOESS residuals < −4 and linear model Cook’s distance > 4/Ncells were called as low-complexity outliers and discarded. Next, we selected cells with cutoffs of >1000 nUMIs, > 200 unique genes, < 25% mitochondrial UMIs, and < 25% hemoglobin-related UMIs. We next used data-driven filtration based on median absolute deviation: an upper cutoff of median + median absolution deviation * 2.5 for cutoffs of mitochondrial content, and a lower cutoff of median – median absolute deviation * 2.5 for nUMIs. The final cutoffs for each of the filtering steps were described in **Supplementary Table 1**. Then, using Seurat (v4.4.0) we clustered each sample using the sctransform workflow, using choosing hyperparamters of 3000 features (default for scTransform), 30 principal components (PCs) and Louvain resolution of 0.1 and all other default parameters^58,59^. Finally, using these clusters, we applied DoubletFinder (v2.0.2) with option “sct” = T, and other parameters kept as default^60^ to exclude doublets.

#### Integration, clustering, and cell marker analysis

For integration and batch correction of scRNA-seq samples, RISC (v1.6) was used^61^. Sample DKO_1 (DJ582M11) was used as the reference sample, PCs 1:30 were used for integration and batch correction, and 10 neighbors were used for Louvain clustering within RISC, while all other parameters were default. Then, Seurat (v5.0.2) was used for marker analysis via the FindAllMarker function with the only.pos parameter set to true. For marker prioritization, a marker score was calculated, consisting of ((pct.1 – pct.2) * acg_log2FoldChange), which emphasizes specificity in marker gene expression.

#### Mutation calling from scRNAseq analysis and malignant cell annotation

For calling single nucleotide variations and clustering based on cell mutational data, Souporcell was used^22^. In most samples, clear distinction of malignant versus stromal clusters was possible using k = 2. For some samples where the distinction was unclear, k =3 was used.

For calling copy number variations (CNVs), InferCNV was used^24^. Macrophages were used as non-mutant reference cells, because these cells were present in all samples. Raw (non-batch corrected) counts were supplied to InferCNV. CNV calling was done separately for non-malignant stromal cells and malignant cells, as InferCNV was unable to run on the entire dataset. A mouse reference annotation GTF file (GenCode VM23) was supplied to InferCNV. For malignant cell analysis, the InferCNV “analysis_mode” parameter was set to “subclusters” to capture intra-tumor heterogeneity. Hidden Markov Model (HMM) gene-level CNV results were used from InferCNV for downstream analysis (InferCNV output filename: “20_HMM_pred.repr_intensitiesHMMi6.leiden.hmm_mode-subclusters.Pnorm_0.5.infercnv_obj”). Malignant and stromal results were read in and harmonized, with macrophage CNV calls excluded from downstream analysis. To study the numbers of genes affected by CNVs for each cell, the number of genes with HMM status not equal to 1 was counted. To study the number of 2X deletions or amplifications per cell, the number of genes with HMM status equal to 0 and greater than or equal to 2 was counted. The HMM matrix was also used for PCA for all cells and for malignant cells only, using the top 1000 most highly variable features for both.

Clusters were annotated to celltypes based on marker analysis and low number of CNVs for immune cells and endothelial cells, using the markers shown in **Figure 1**. Final cell annotations for malignant cells and cancer associated fibroblasts was based on marker analysis, SouporCell clustering and InferCNV results. The fibroblast cluster was annotated as such because its small number of inferred CNVs matched that of immune and endothelial cells, and clustered with the rest of the microenvironment in Souporcell results for most samples. Integrated clustering took precedence over individual sample clustering in cell annotation.

#### Differential expression analysis across TKO, DKOAA and DKO

To take advantage of replicates, a pseudobulk based strategy was used to reduce false positives^62^. For each celltype, UMIs for all cells were summed up by sample for each gene, then gene expression among the three OS models were compared via the EdgeR Likelihood Ratio Test^63^. Gene set enrichment analysis (GSEA) was applied via the FGSEA package for pathway analysis^64,65^. Gene sets including the Hallmarks gene sets were derived from the Molecular Signatures Database via the “Msigdbr” package^66–68^. For network analysis of enriched pathways, aPEAR (v1.0.0 from CRAN) was used^69^.

#### Subclustering and differential compositional abundance analysis

For subclustering of individual celltypes, RISC batch-corrected data from each cell type was subsetted and Seurat (v4.4.0) was used to identify the top 2000 most highly variable genes and scale the expression of these genes before PCA. For most celltypes, PCs 1:30 and Louvain resolution of 0.5 was used. In some celltypes, poor overlap between samples was observed and marker analysis did not reveal clear cluster markers, so these values were adjusted. In particular, for sub-clustering of malignant cells, PCs 1:10 and resolution 0.1 were used, while for macrophages, PCs 1:30 and resolution 0.7 were used. Markers were derived for each subcluster using Seurat function FindAllMarkers with the parameter “only.pos” set to True.

Compositional analysis was also applied in a pseudobulk-aware manner to reduce false positives^70^. For compositional analysis between the three OS models, the Propeller test was used with option “transform” set to “asin” for arcsine square root transformation, as recommended by benchmarking analysis^70,71^.

#### Differential CNV analysis across TKO, DKOAA and DKO

To compare patterns of CNVs across the three OS models, we devised two novel replicate-aware methods. Both were based on the gene-level Hidden Markov Model (HMM) from InferCNV (see above). In the first, which we referred to as the “differential CNVs mean” or “DiffAmps_mean” approach, we took the mean CNV state of all cells for each sample, resulting in a matrix of CNV gene by sample, where values were the mean CNV state of all cells from that sample. Then, for each gene, the difference between OS models were compared in a replicate-aware manner via

Wilcoxon test. In the second method, which we referred to as the “differential CNVs count” or “DiffAmps_count” approach, we separately considered amplifications and deletions. For each gene, the number of cells with HMM state of 0 (2X deletion) or 2 or more (2X or higher amplification) was counted. Then, for each sample, the proportion of cells with or without 2X deletion or amplification was calculated. For each sample, this produced a matrix of gene by sample, where each value was the proportion of cells with extreme (2X) CNV. Because this was equivalent to compositional abundance analysis, we then transformed the proportions using the arcsine square root transformation and compared via t-test, similar to Propeller analysis (see above). For both methods, p < 0.1 is used, on the basis of this being a secondary, exploratory analysis. However, to increase biological relevance, only genes that were also significantly overexpressed by differential expression analysis were considered for downstream analysis and literature search.

#### Signature score analysis, cell cycle analysis, and Myc proportion analysis

To study expression of a group of genes, the previously described module score method was used to compute signature score for the gene group^72^. This method is implemented in Seurat in the “AddModuleScore” function. To study cell cycle phase, we used the related “CellCycleScoring” with genes derived also from Seurat lists of S phase and G2/M phase genes. To study *Myc* responder cells, we used all genes from the “Hallmark *Myc* Targets V1” gene set to calculate *Myc* target signature scores among malignant OS cells. Next, we defined *Myc*-Responder High cells as those with a score above the median value determined for all malignant cells in all samples. Then, we compared proportions of *Myc*-Responder High cells between OS models in a replicate-aware manner using the Propeller test with the “asin” transformation, as described see above.

#### Human OS scRNA-seq data analysis

To compare murine OS with human patient OS, we downloaded the data from two studies that applied scRNA-seq to human patient OS tumors^20,21^. Data was downloaded from the GEO (accessions numbers: GSE152048 and GSE162454). Datasets were integrated, batch-corrected, and clustered using RISC (v1.6) as described above for the mouse OS data. Cell annotations were not shared publicly by either study, so we performed cluster marker analysis using Seurat (v5.0.2) FindAllMarkers function with only.pos set to true. For immune cells, clusters were annotated using marker genes as shown in **Figure 6**. To identify malignant OS cells and non-malignant, non-immune stromal cells, InferCNV was applied, using macrophages as reference cells. InferCNV was applied to each sample individually, then HMM results were harmonized across samples and macrophage CNV calls were not used in downstream analysis. Malignant clusters were called based on presence of CNVs and markers, while fibroblasts and pericytes were called based on their lack of CNVs and markers.

To study the fidelity of murine OS to human tumors, markers from murine cell types were used. Using the package “biomaRt” (v2.54.1)^73^ (https://feb2021.archive.ensembl.org/), human orthologs were found and only markers with a single unambiguous ortholog were retained. Using these ortholog lists, signature scores for each murine celltype were calculated in the human OS data. The average score for each human celltype was calculated, and then a correlation matrix was computed and used for hierarchical clustering.

#### NCI TARGET OS survival analysis

We downloaded clinical and transcriptomic data from the NCI TARGET OS cohort using the TARGET data portal^74,75^. After downloading the data, we filtered for samples which we could harmonize with clinical and survival data, and removed samples with abnormally low TPM values, leaving 83 patients. The R package biomaRt was used to find orthologous pairings between mouse and human genes, while genes that could not be unambiguously mapped were removed^73^. To calculate expression signature scores for the murine OS marker lists, the TKO and DKOAA upregulated interferon scores, and the TKO upregulated myogenesis score, we used the module score method^72^. For survival analysis with the module scores, we used the survival (v3.4.0) and survminer (v0.4.9) packages in R for Kaplan-Meier (KM) analysis. For KM we dichotomized the module scores at the median value to compare survival in high versus low with KM and log-rank tests.

#### Label transfer from bone atlas and inference of OS pathologic subtype

To infer pathologic subtypes (osteoblastic, chondroblastic, fibroblastic), a scRNA-seq dataset of non-malignant murine bone cells was used^48^. Counts and metadata were downloaded from the Broad single cell portal (Accession: SCP361). Cell annotations were simplified to combine cell clusters together by cell type (ie, Fibro 1,2,3,4, and 5 were combined to “Fibro”). Data was downloaded and processed with the Seurat “sctransform” function. Then, Seurat label transfer was used. For each murine OS sample, malignant cells were selected, then the function FindTransferAnchors was used with healthy bone as the reference and malignant OS as the query, PCs 1:30, and normalization method set to “SCT”. Then, TransferData was used to acquire inferred celltypes and scores. To call a whole tumor as a pathogenic subtype, the transferred celltype with the highest proportion among malignant cells was used as the label. To compare proportions of microenvironment cell types between osteo and fibro subtypes, where multiple samples were detected, we used the propeller test.

#### Metastatic status evaluation of lung

Mice of each model with observable limb tumors that met euthanasia criteria were continuously included in the study. After euthanasia, whole lung tissue was collected from 19 DKO mice, 17 DKOAA mice, and 38 TKO mice. The harvested lung tissue was subsequently rinsed in cold PBS and fixed with formalin. Paraffin blocks were created, and serial sectioning was performed at consistent intervals, with 10-30 intervals per lung determined based on lung size. A total of 480 intervals from DKO mice, 400 from DKOAA mice, and 515 from TKO mice were obtained. Each interval underwent H&E staining and was examined for metastatic lesions by two practicing clinicians. The proportion of lungs and intervals affected by metastatic lesions was calculated and compared among the various groups via Fisher exact test.

## Data Availability

The data generated in this study are publicly available in the Gene Expression Omnibus (GEO accession number: GSE262743).

For exploratory data analysis, we provide two web-based Shiny apps, generated using the ShinyCell method^76^. For murine data, the app is available here: https://scviewer.shinyapps.io/mouseos_scrnaseq_tko_dkoaa_dko/. For integrated human data, the app is available here: https://scviewer.shinyapps.io/humanos_scrnaseq/.

## Authors’ Contributions

**A. Ferrena**: Data curation, software, formal analysis, investigation, visualization, methodology, writing–original draft, writing–review and editing. **J. Wang**: Resources, data curation, formal analysis, investigation, methodology, writing–review and editing. **R. Zhang**: Resources, data curation, validation, visualization, methodology, writing–review and editing. **X.Z. Zheng**: Software, formal analysis, visualization, writing–review and editing. **B. Göker**: Resources, methodology. **H. Borjihan**: Resources, methodology. **S.S. Chae**: Resources, methodology. **Y. Lo**. Methodology. **H. Zhao**: Resources, writing–review and editing. **E.L. Schwartz**: Resources, writing–review and editing. **D. Loeb**: Resources, writing–review and editing. **R. Yang**: Resources, methodology. **D.S. Geller**: Resources, methodology. **D. Zheng**: Conceptualization, supervision, funding acquisition, writing–original draft, project administration, writing–review and editing. **B. H. Hoang**: Conceptualization, resources, supervision, funding acquisition, writing–original draft, project administration, writing–review and editing.

## Acknowledgments

This study is supported by the National Institutes of Health (NIH/NCI; R01CA255643 to B.H. Hoang, D. Zheng, Y. Lo, H. Zhao, E. Schwartz), Sarcoma Strong (to B.H. Hoang), and the Arnold and Madaleine Penner Endowment fund (to B.H. Hoang). A. Ferrena is supported by the PhD in Clinical Investigation program at Albert Einstein College of Medicine under NIH/National Center for Advancing Translational Science (NCATS) Einstein-Montefiore CTSA Grant Number (UL1TR001073) and NIH TG number (TL1TR0022557). J. Wang would like to acknowledge the supports of National Natural Science Foundation of China (82103223 to J. Wang), Peking University People’s Hospital Scientific Research Development Funds (RDX2022–01 to J. Wang), and Peking University Clinical Scientist Training Program (BMU2023PYJH015 to J. Wang).

