## Supplemental Figures for "Comprehensive single cell transcriptomics analysis of murine osteosarcoma uncovers *Skp2* function in metastasis, genomic instability and immune activation and reveals additional target pathways"

### Supplementary Materials

**Supplementary Figures 1-21**

**Supplementary Tables 1-9 (in an Excel file)**

**Supplementary Figure 1: Clustering and markers of transgenic OS tumors.** A: Louvain clusters of integrated scRNA-seq data (from RISC). B: Violin plot of number of genes affected by "extreme CNVs" (>=2x deletion or amplification). C: Canonical and data-driven markers plotted for the clusters in A; the same markers as Fig 1C are shown. D: Data-driven markers for each cluster. E: Data-driven markers for each cell type after merging clusters for the same cell type. F: Correlation matrix and hierarchical clustering of cell types across samples, based on average gene expression in each cell type per genotype. G. UMAP of Osx (*Sp7*) expression pattern.

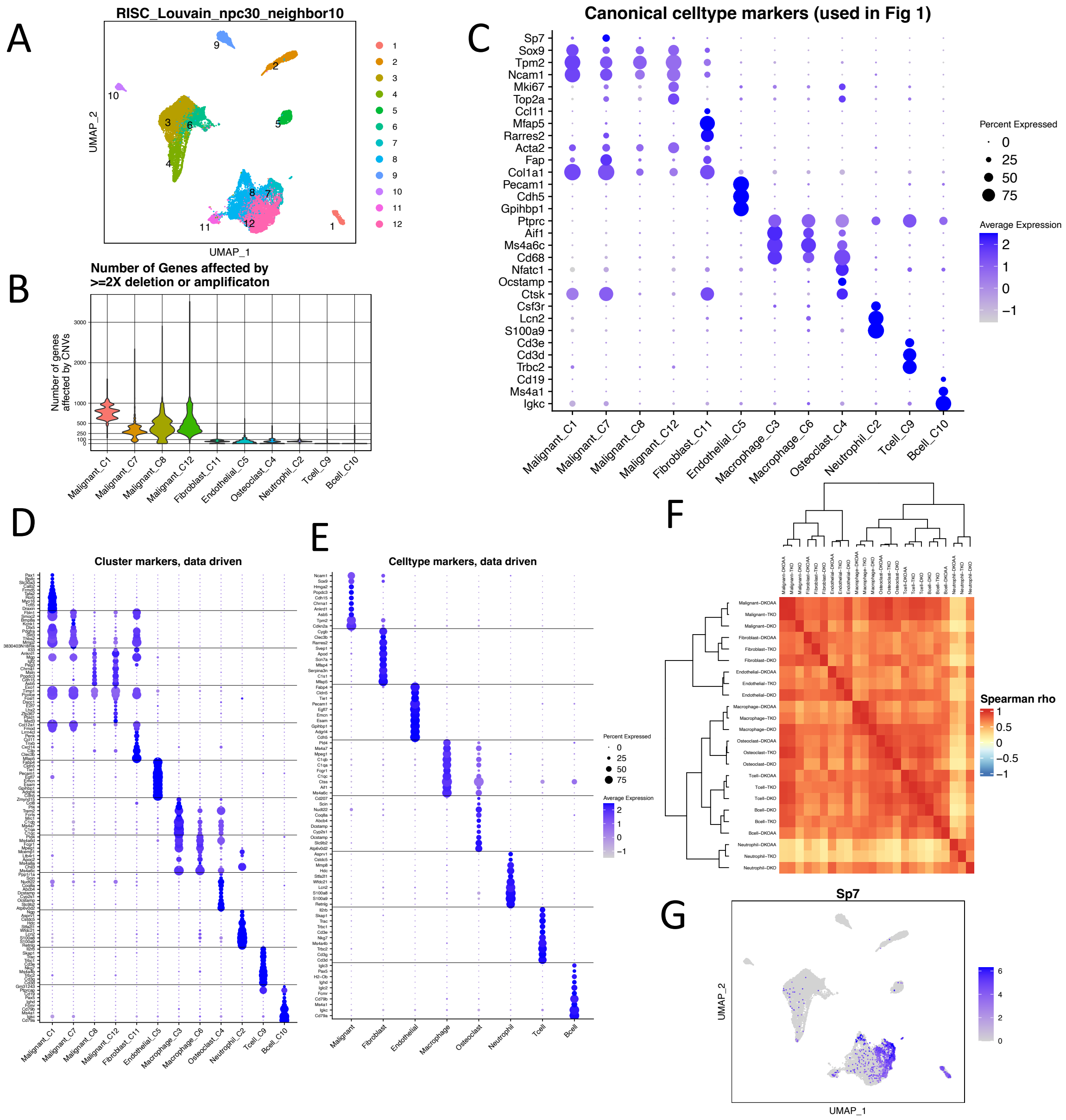

**Supplementary Figure 2: Souporcell results.** For each sample, two UMAP plots are shown: UMAP derived from Seurat based on transcriptional data, colored by final integrated cell type (left) or by each sample's Souporcell clustering results (right). The K parameter for Souporcell clustering is also shown. K = 2 was used for all samples except those where a clear malignant cluster could not be distinguished. For DKO_2, DKO_3 and DKOAA_2, K was set to 3.

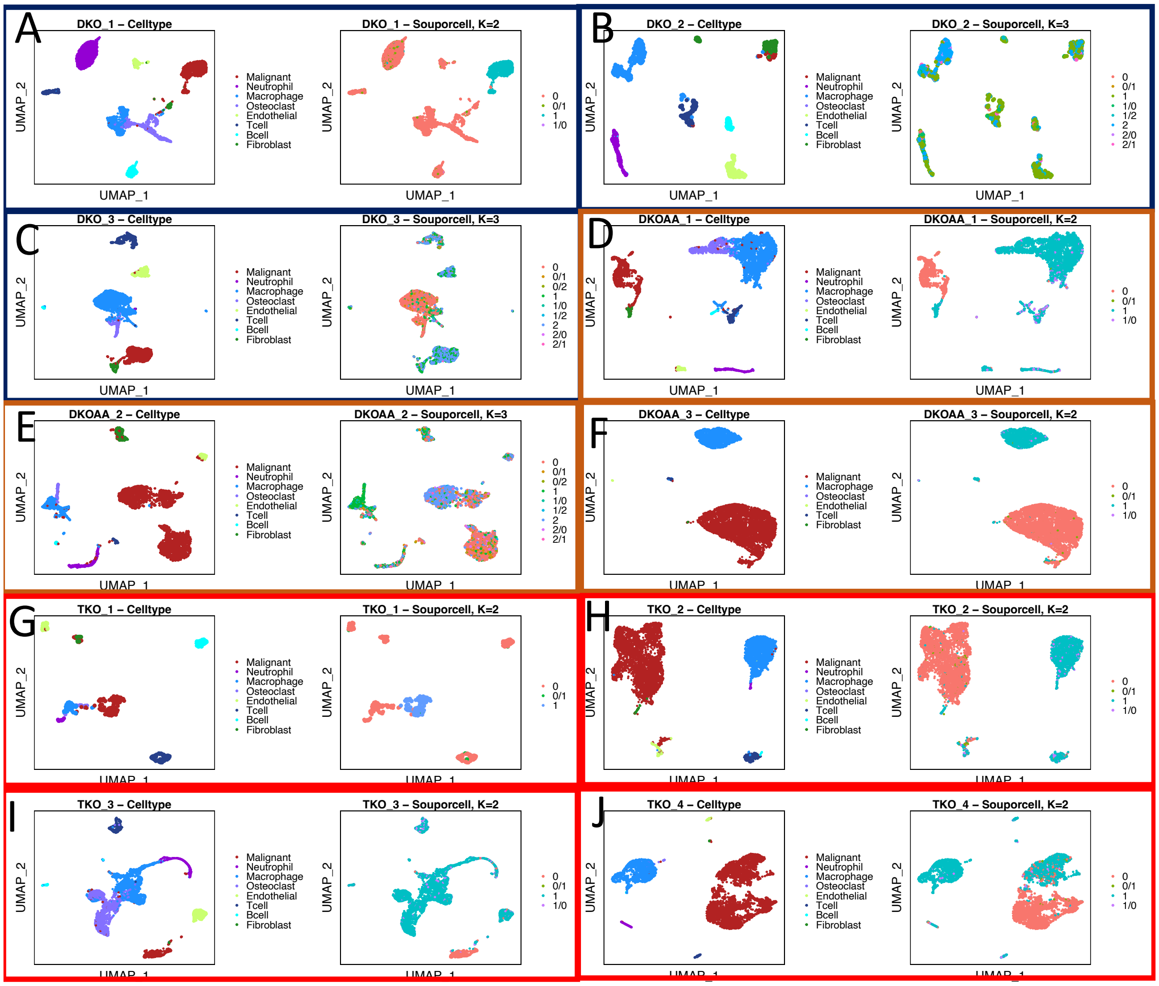

**Supplementary Figure 3: side by side comparison of InferCNV results for stromal cells (top) and malignant cells (bottom).** Macrophages were used as reference celltypes in both. A: InferCNV results for stromal cells from all samples. B: InferCNV results for malignant cells from all samples.

A

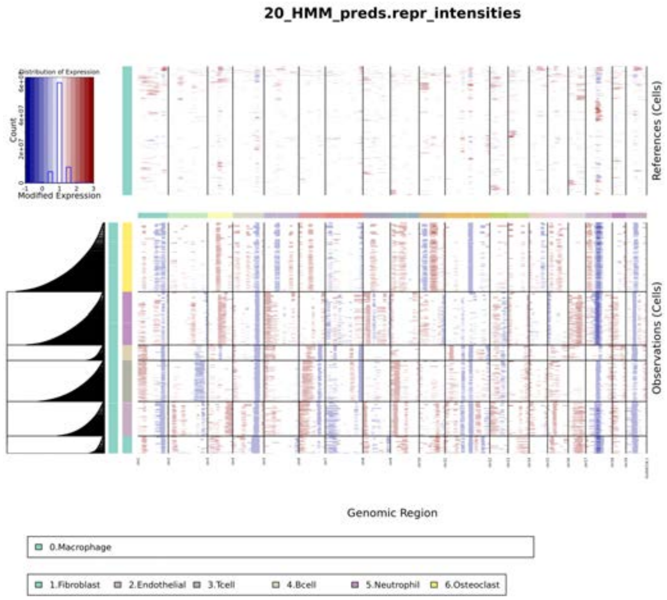


B

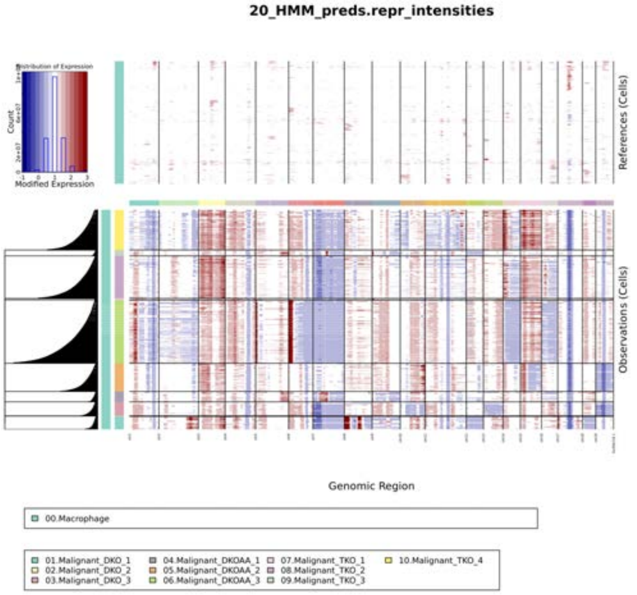

**Supplementary Figure 4. Leading edge genes from GSEA showing downregulation of invasive phenotypes in TKO and DKOAA relative to DKO.** The intersects of leading edge genes from both TKO and DKOAA versus DKO enrichments are shown. A: Leading edge genes for Hallmark Epithelial-Mesenchymal Transition gene set. B: Leading edge genes for Reactome Extracellular Matrix Organization gene set. C: Leading edge genes for Gene Ontology Biological Process Cell Migration gene set.

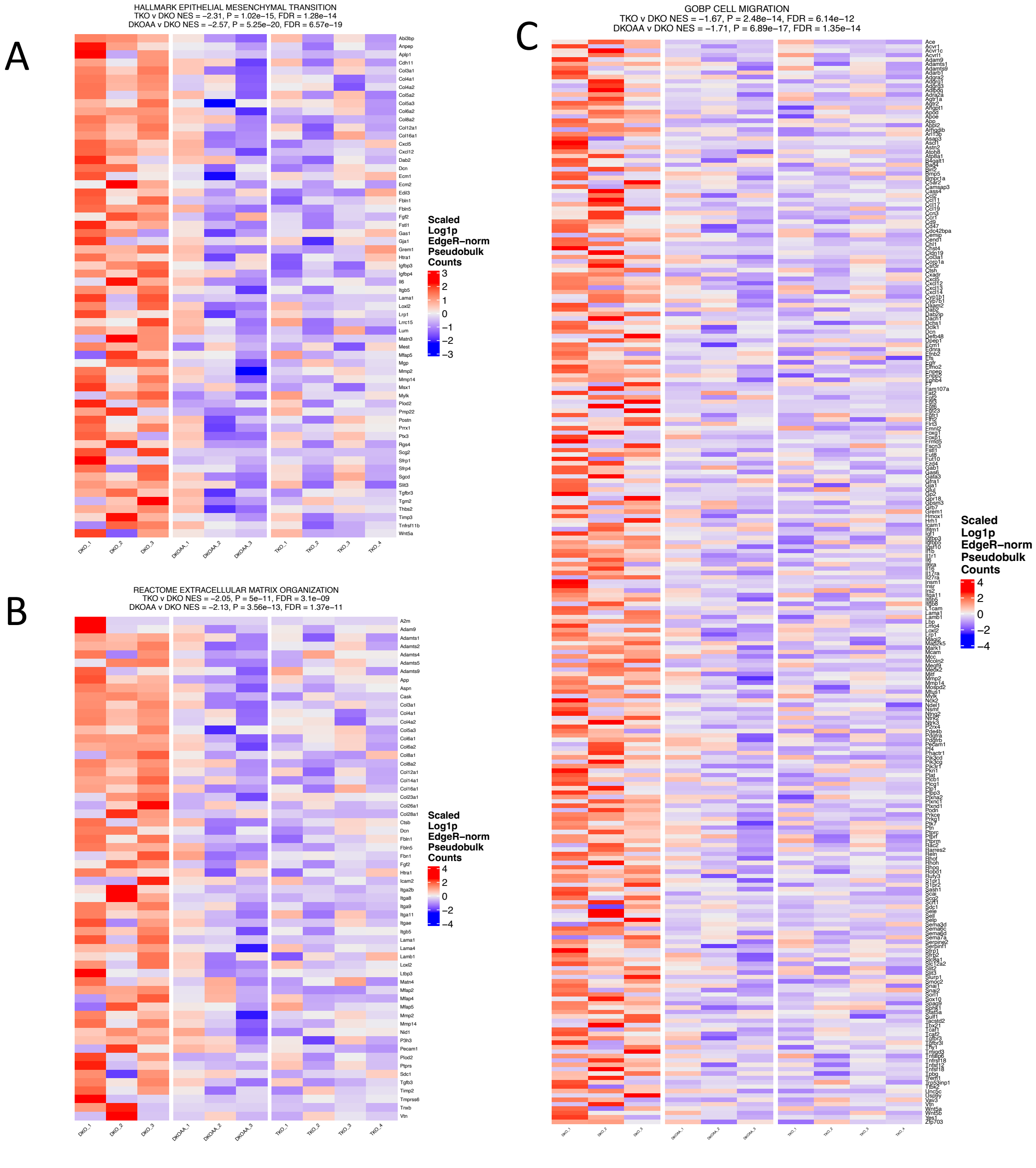

**Supplementary figure 5: Differential pathway gene set enrichment analysis among TKO vs DKO in all cell types, visualized as networks via aPEAR.** Results are shown for each celltype.

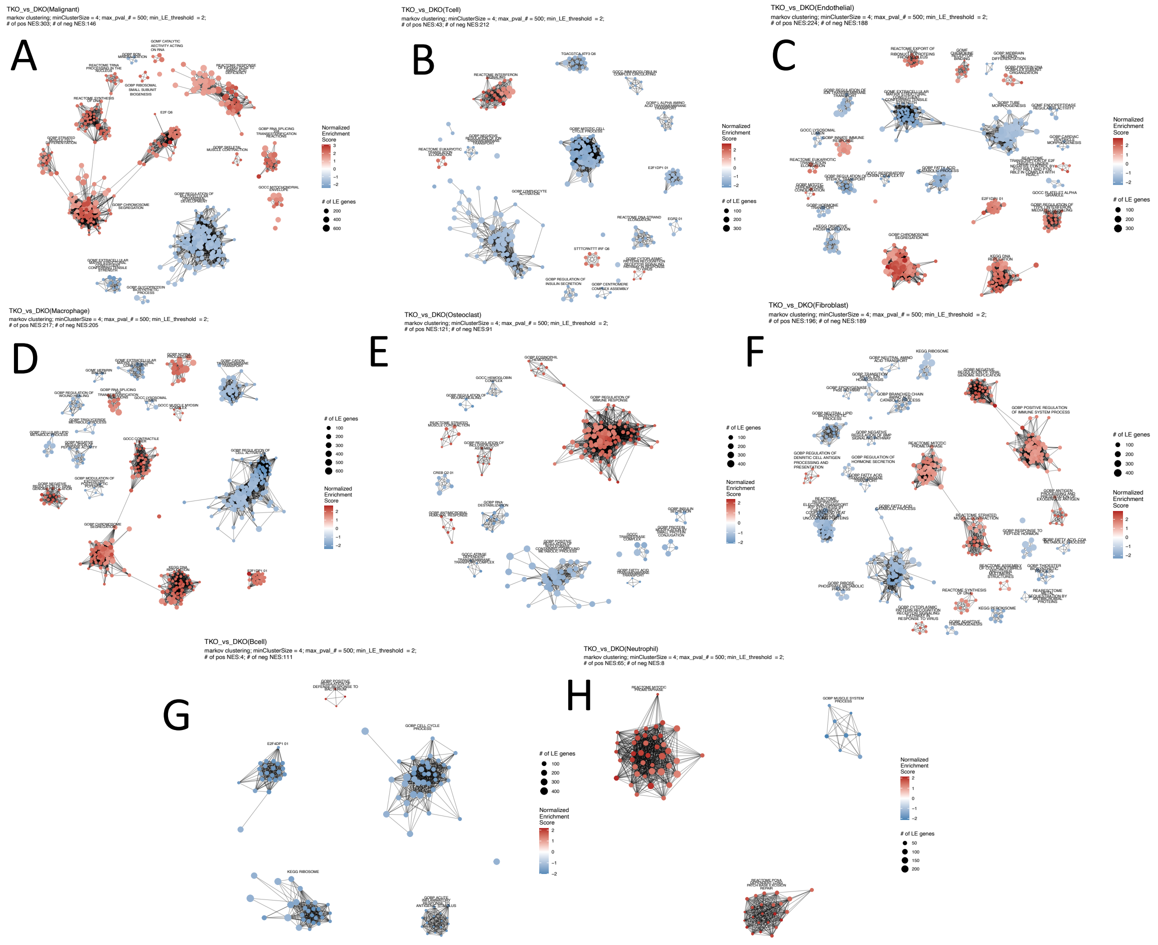

**Supplementary figure 6: Differential pathway gene set enrichment analysis among DKOAA vs DKO in all cell types, visualized as networks via aPEAR.** Results are shown for each celltype.

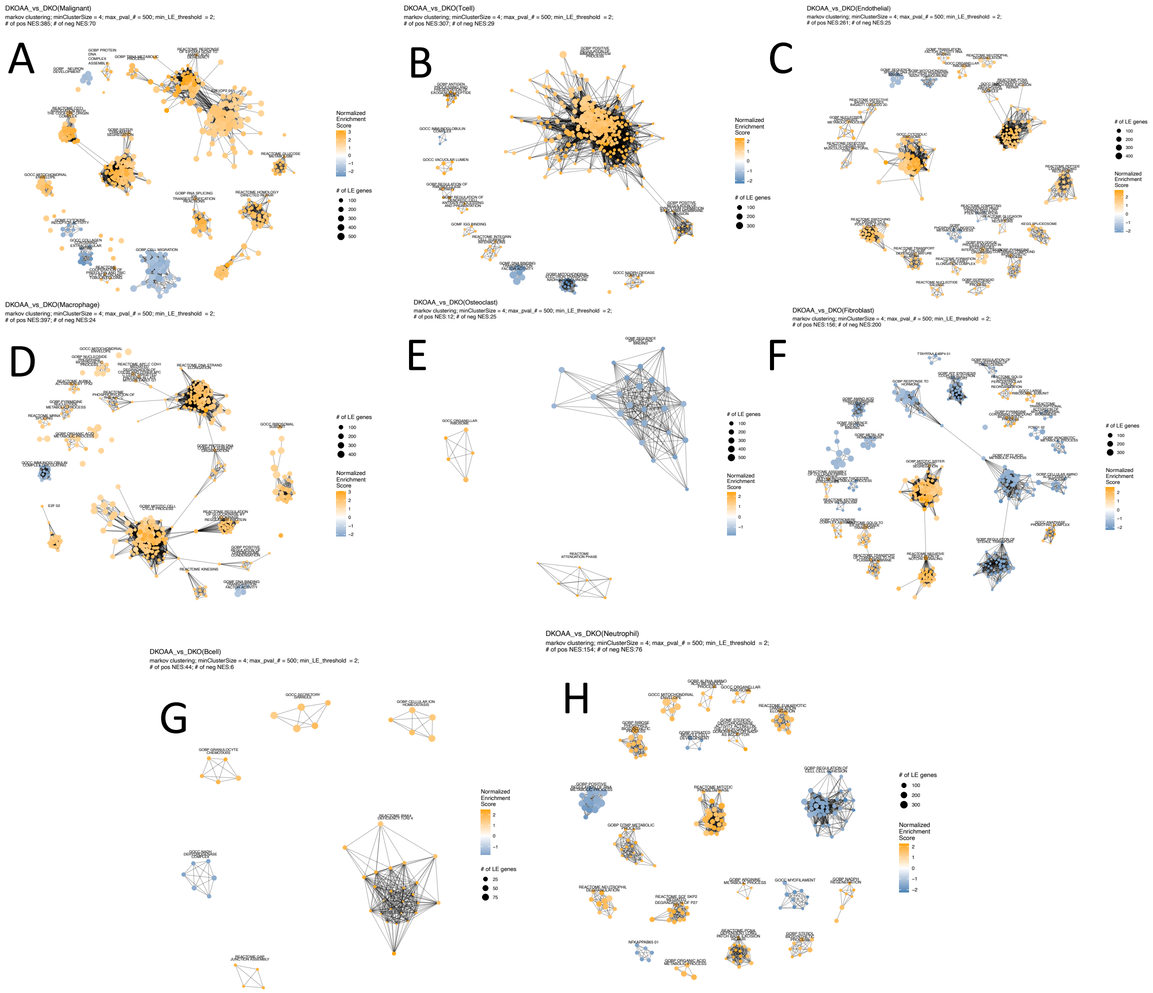

**Supplementary Figure 7: GSEA of DKOAA and leading edge genes of interferon alpha and gamma response.** A: Dot plot showing GSEA results of Hallmarks gene sets in DKOAA cell types relative to DKO. B-F: Heatmaps showing leading edge genes of Hallmark IFN-alpha and IFN-gamma response gene sets in the cell types indicated. The union of IFN-gamma and IFN-alpha leading edges is shown. Additionally, the union of TKO and DKOAA leading edges from these gene sets is shown for T cells and Osteoclasts, while for other celltypes, only the significant TKO enrichment's leading edge is shown.

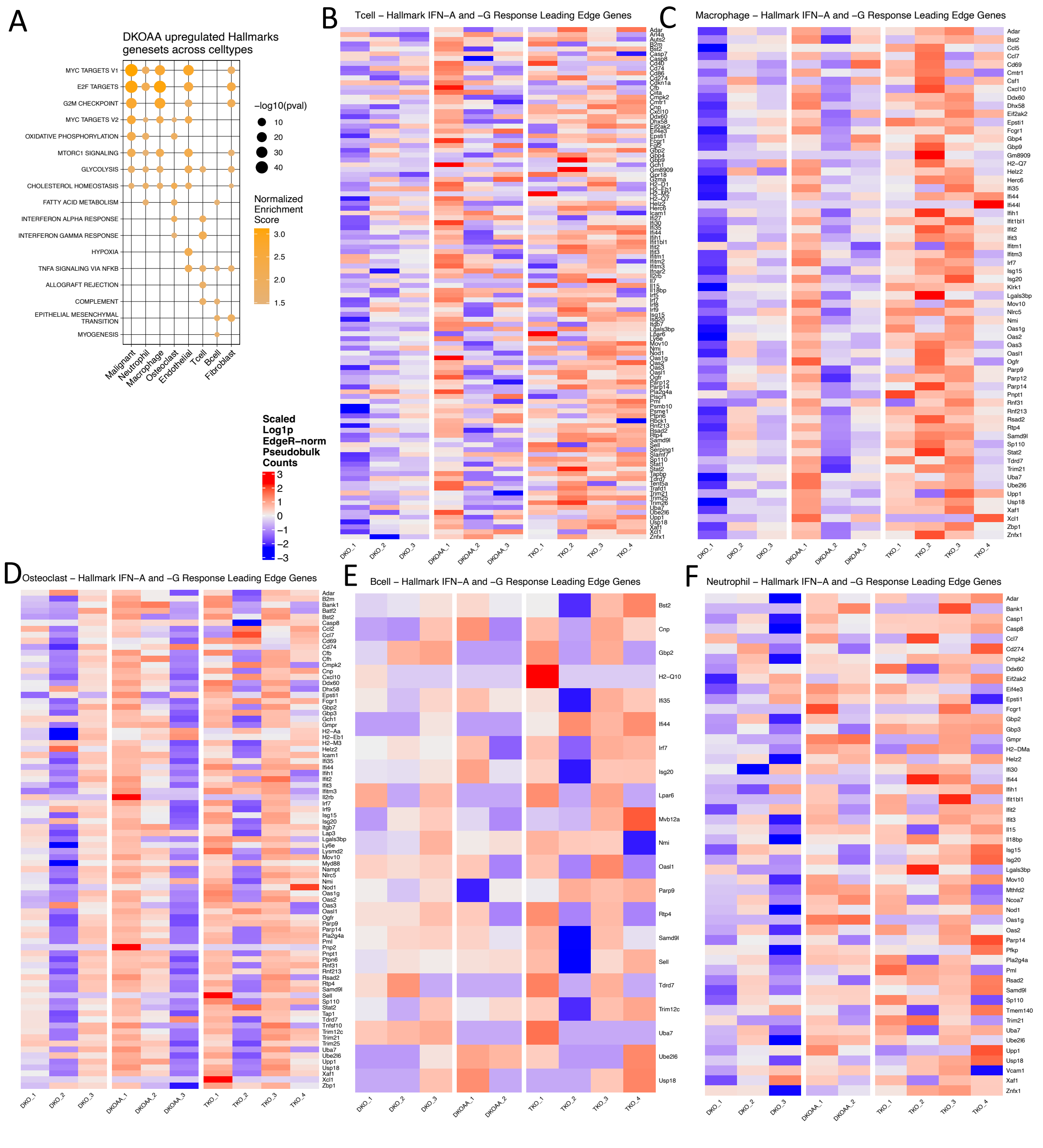

**Supplementary Figure 8: GSEA of genesets related to cellular stress response.** A,B: Dot plot showing GSEA results for gene sets significantly differentially expressed in TKO and DKOAA versus DKO, respectively. C – F: Heatmaps showing leading edge genes from significantly enriched gene sets of all cells. Star callouts are used to indicate that a significant enrichment was observed.

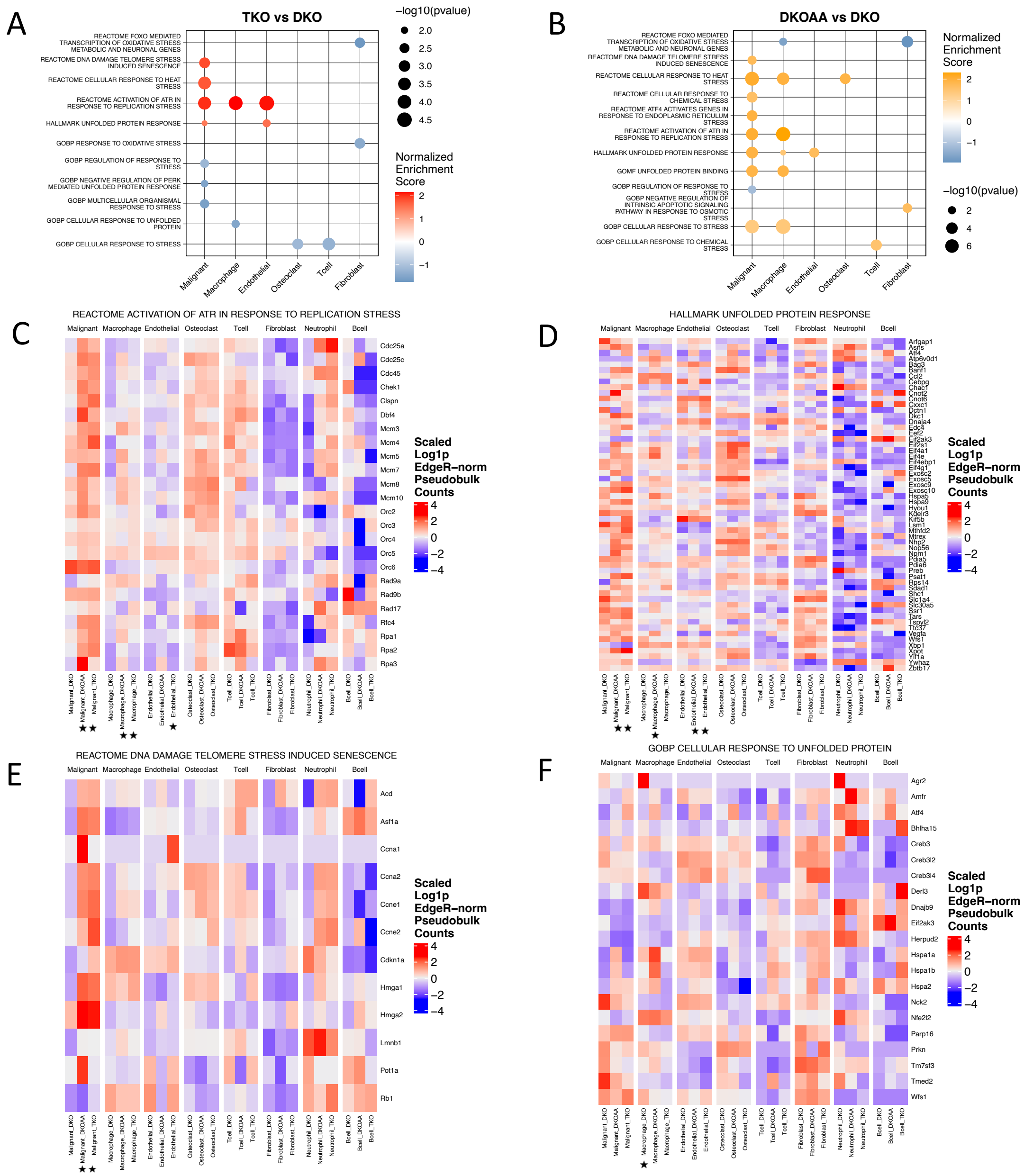

**Supplementary Figure 9: Analysis of Interferon related genes.** A: UMAP plots showing expression patterns of interferon genes. B,C: Dotplot and heatmap showing expression differences of interferon related genes across OS models, stratified by celltype.

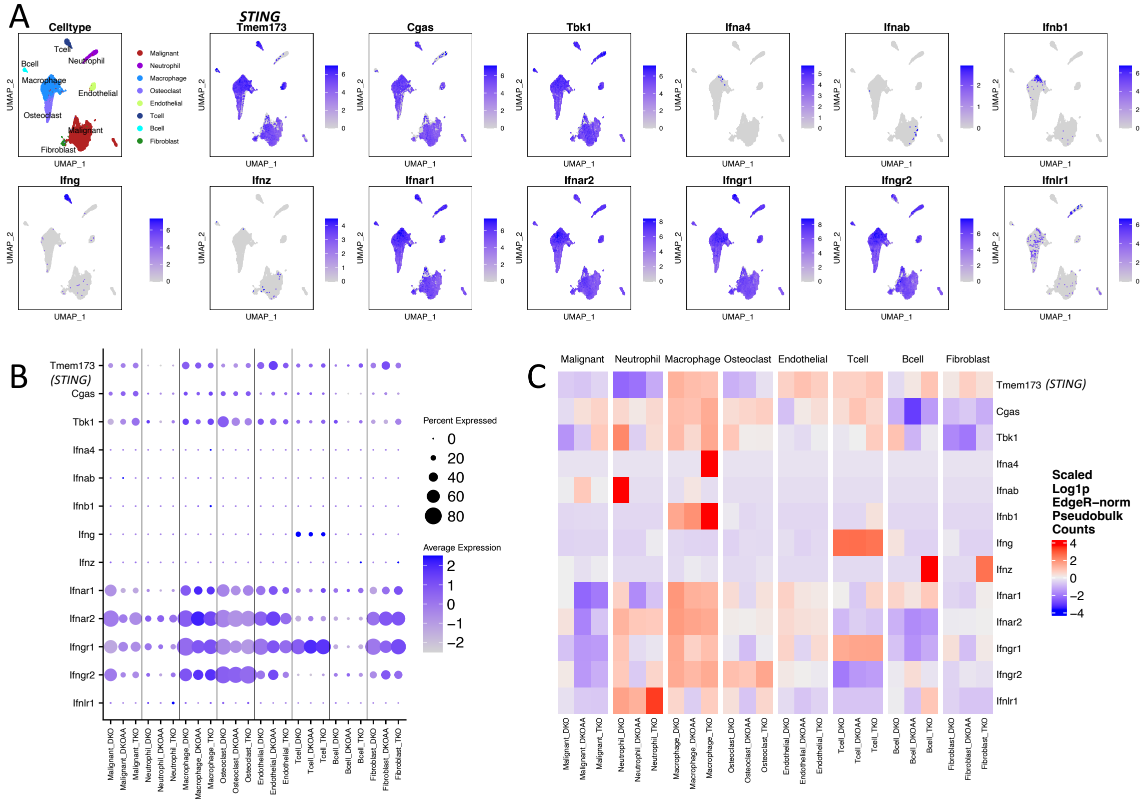

**Supplementary Figure 10: CellChat analysis of cell signaling differences between OS tumors.** A: Differential signaling strength among cell types in TKO vs DKO. Red = increased in TKO. B: Differential signaling strength among cell types in DKOAA vs DKO. C-E: Outgoing signaling pathways detected across TKO, DKOAA, and DKO. F-H: Incoming signaling pathways detected across TKO, DKOAA and DKO.

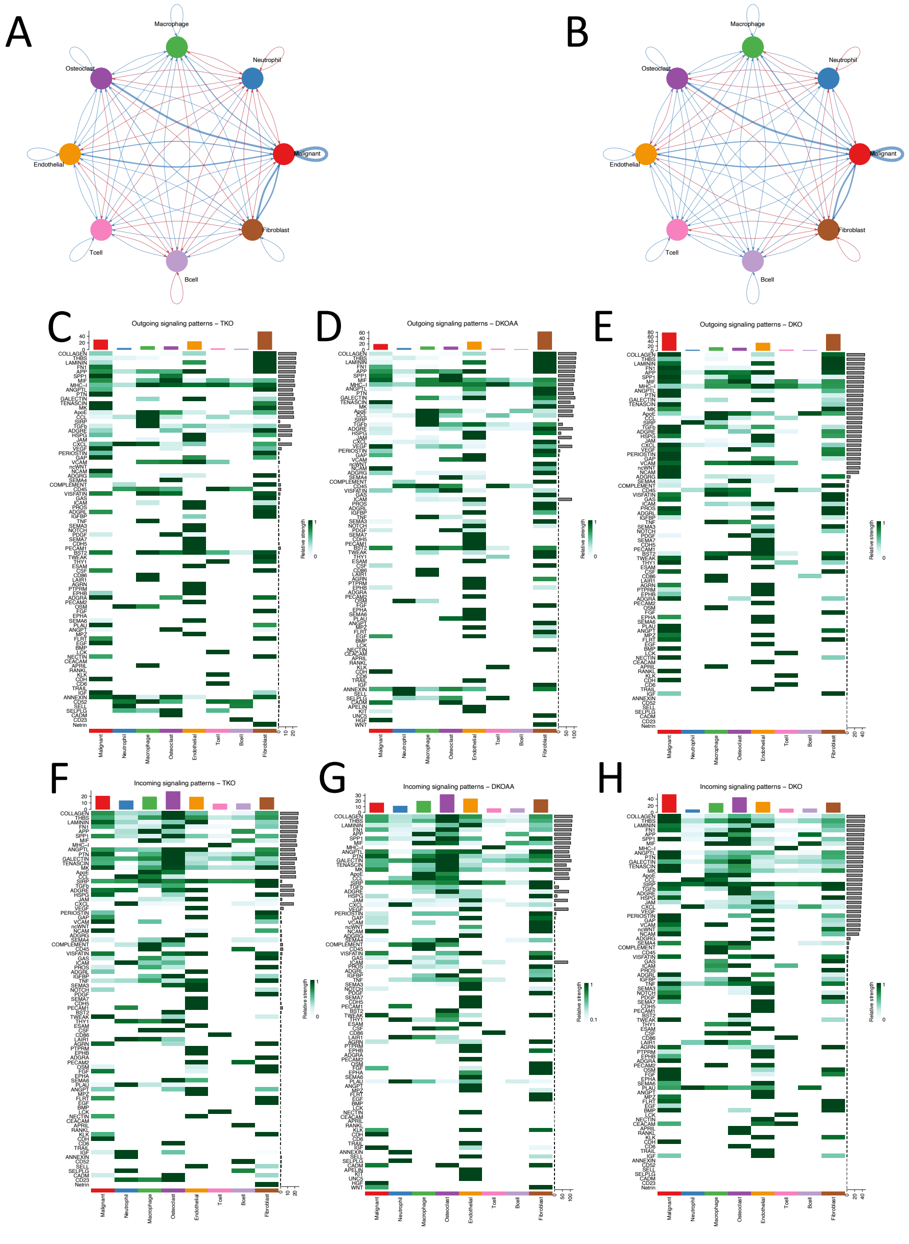

**Supplementary Figure 11: Subclustering of individual cell types.** Two UMAP plots are shown for each cell type: on the left, colored by Louvain sub-clusters; on the right, colored by OS models.

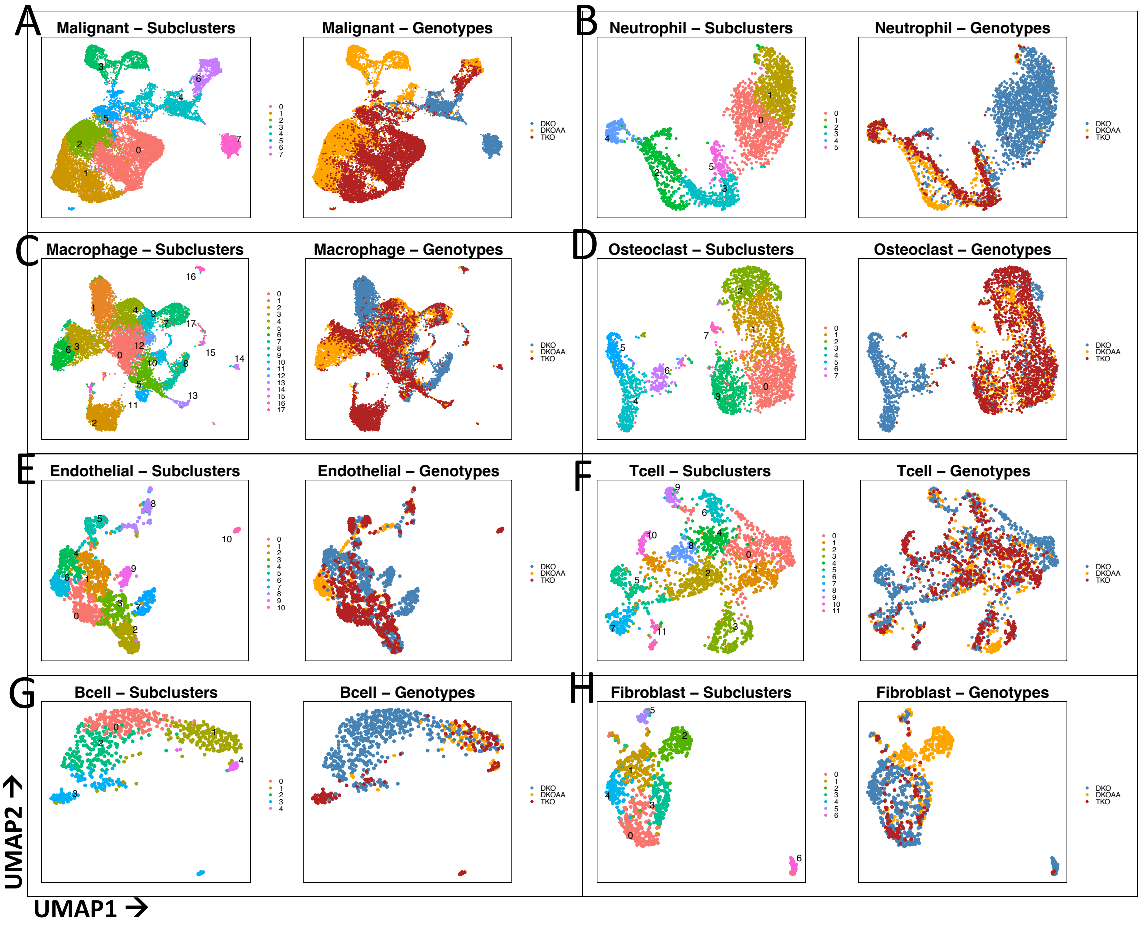

**Supplementary Figure 12: Marker analysis for subclusters of individual cell types.** The top 5 markers are shown for each sub-cluster.

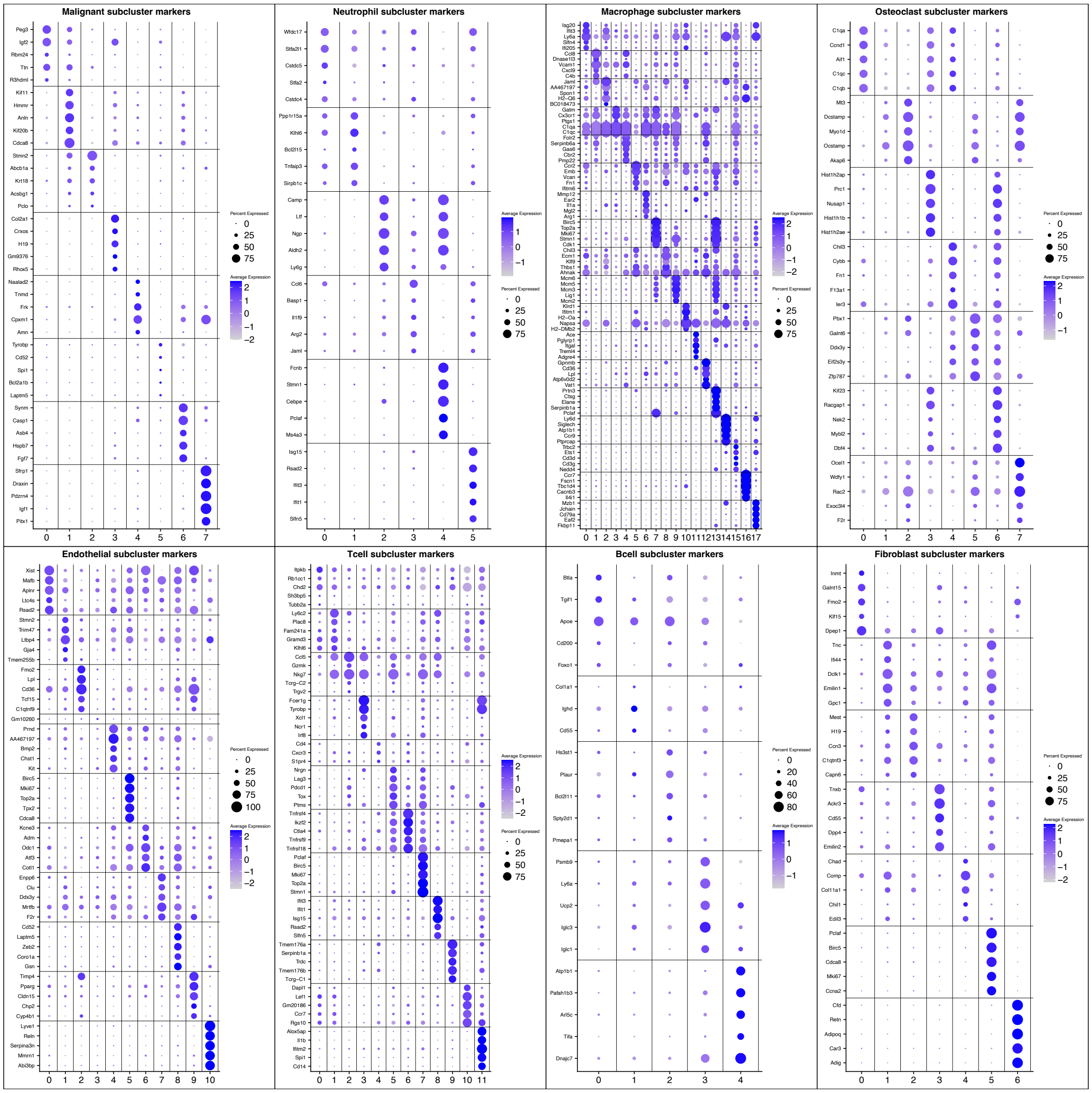

**Supplementary Figure 13: Heatmaps showing proportions of subclusters for cell types containing at least one subcluster with a significant difference of proportion among OS models.** Significant differences ($p < 0.05$) are denoted by cluster labels with asterisks. Propeller test of proportions from the Speckle package was used for statistical testing.

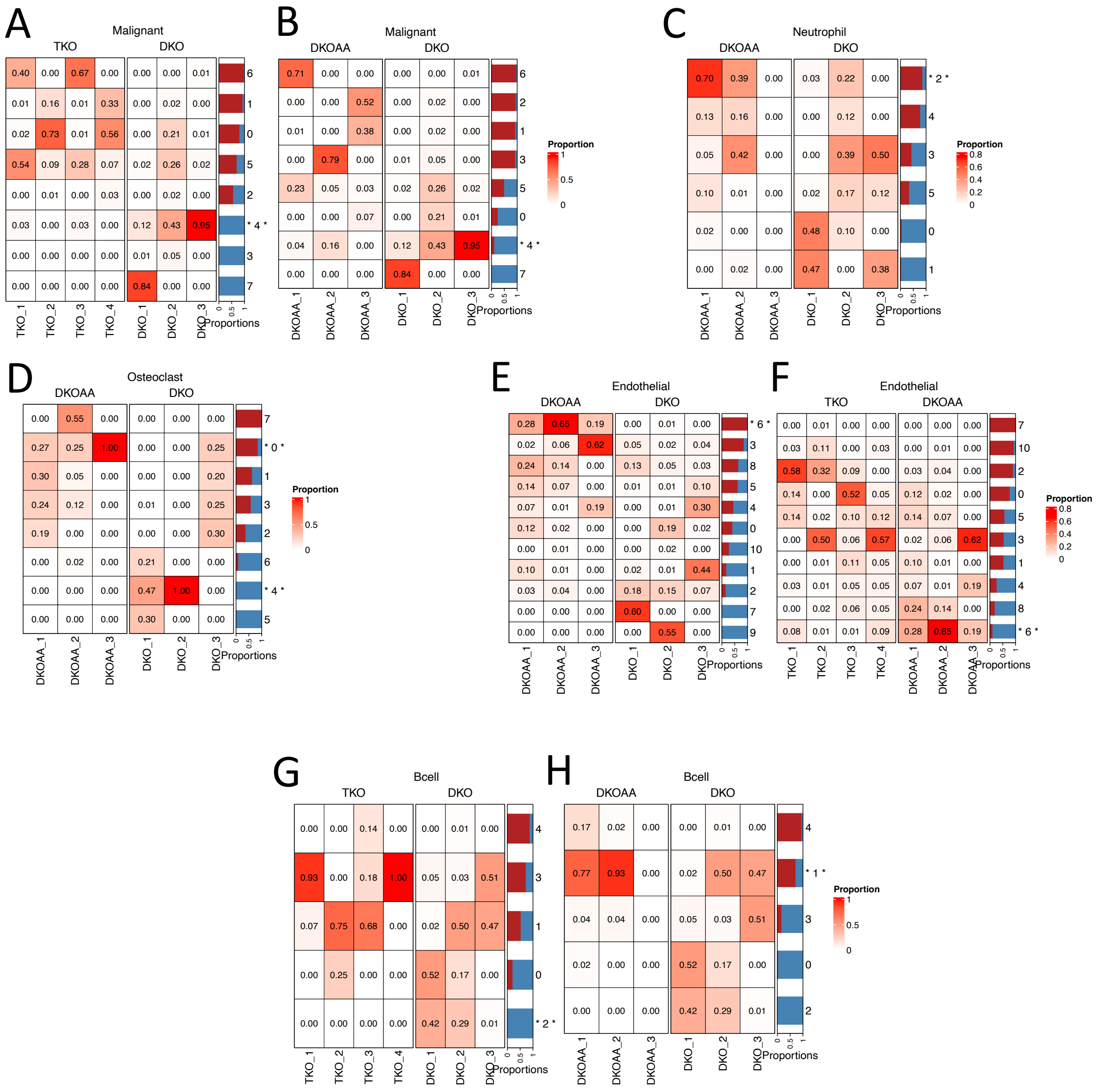

**Supplementary Figure 14: E2f and Myc upregulation in TKO and DKOAA malignant cells.** A: E2f and *Myc* family gene expression patterns across models for malignant cells. B: E2f and *Myc* family gene expression across samples for malignant cells. C: E2f and *Myc* transcriptional regulon scores computed by SCENIC. These are the only transcription factors from the genes shown in A and B for which SCENIC scores were available. D: UMAP of celltypes colored by cell cycle phase. E,F: Violin plots of S phase and G2/M phase scores, respectively. G-K: Heatmaps for the expression patterns of regulon genes from SCENIC for the transcription factors in panel C among malignant cells from TKO, DKOAA and DKO. L: Enrichment and leading-edge genes for the Reactome Apoptosis gene set among malignant cells from TKO, DKOAA and DKO. M: Violin plot of the *Myc* signature score calculated in all cells.

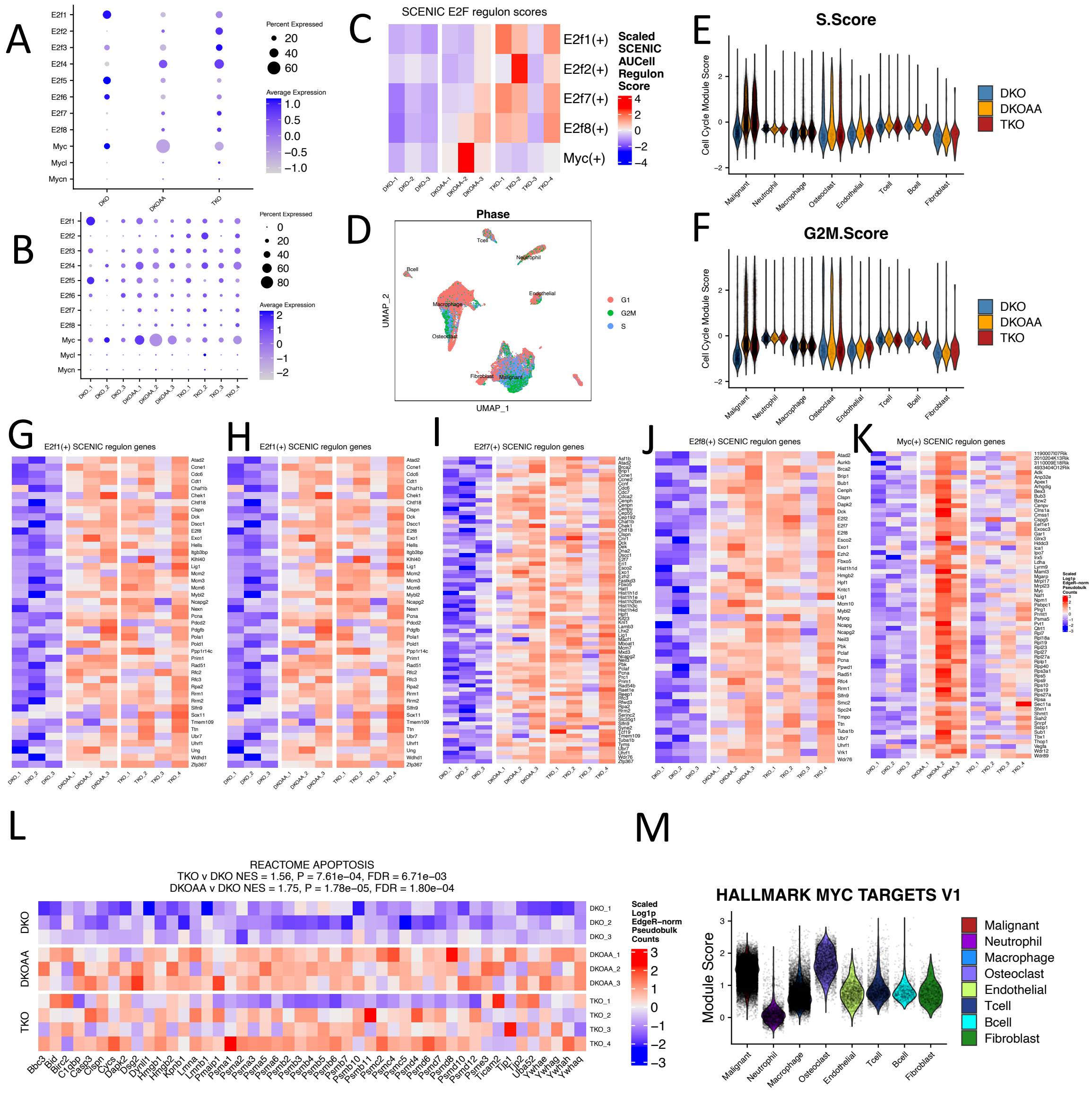

**Supplementary Figure 15: Clustering and markers of integrated human OS scRNA-seq data.** A: Louvain clusters of integrated human scRNA-seq data (from RISC). B: Violin plot of number of genes affected by “extreme CNVs” (>=2x deletion or amplification). C: Canonical and data-driven markers plotted at the cluster level; the same markers as Fig 6D are shown. D: Data-driven markers for each cluster. E: Data-driven markers for each celltype after merging clusters of the same cell type.

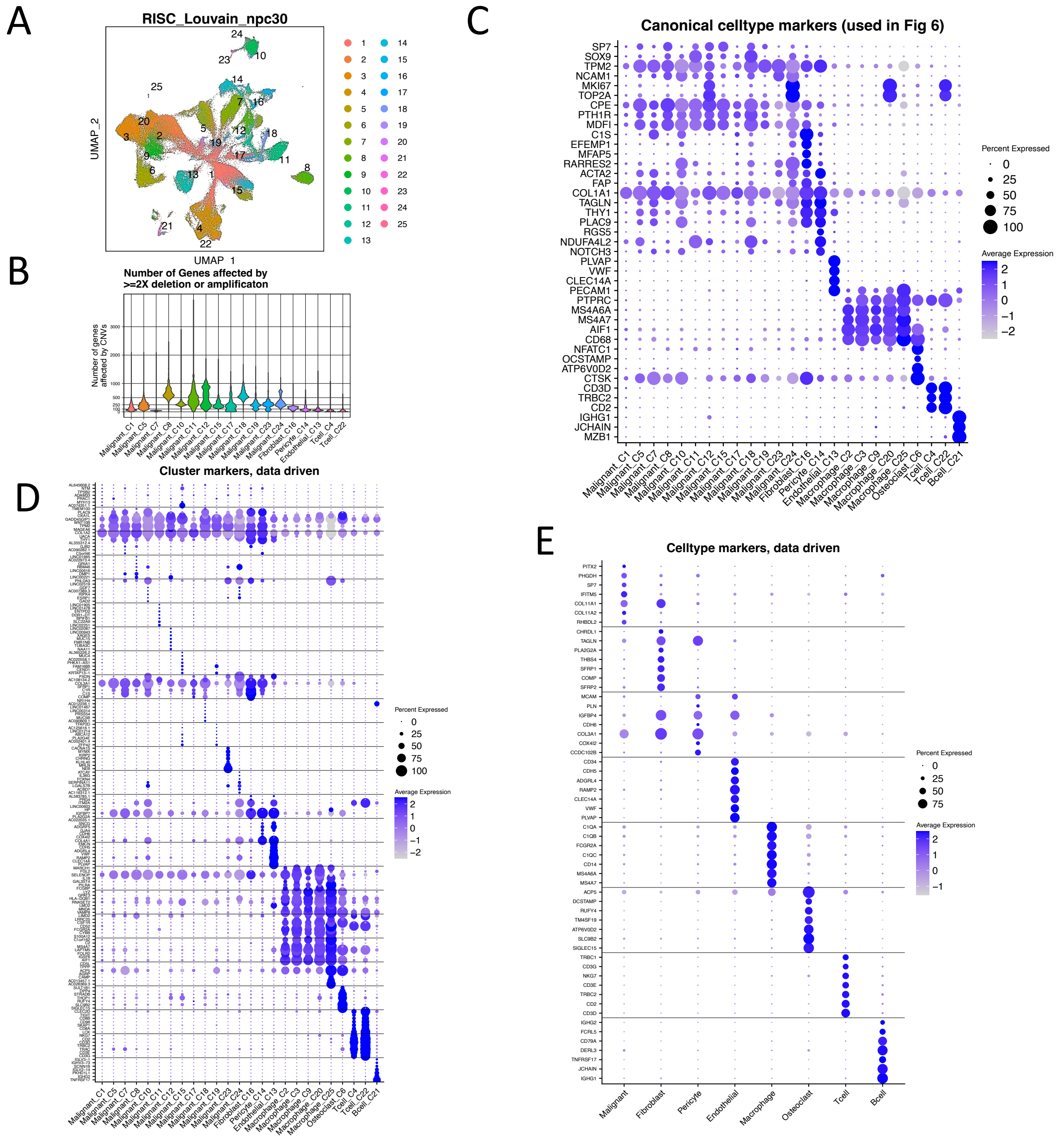

**Supplementary Figure 16. Simplification of bone cell atlas annotations for usage in label transfer.** A: UMAP of atlas data, colored by published clusters. B: UMAP colored by joined, simplified celltypes. C: Markers of simplified celltypes plotted in published clusters. D: Markers of simplified celltypes plotted after joining clusters to simplified, cell-type annotations.

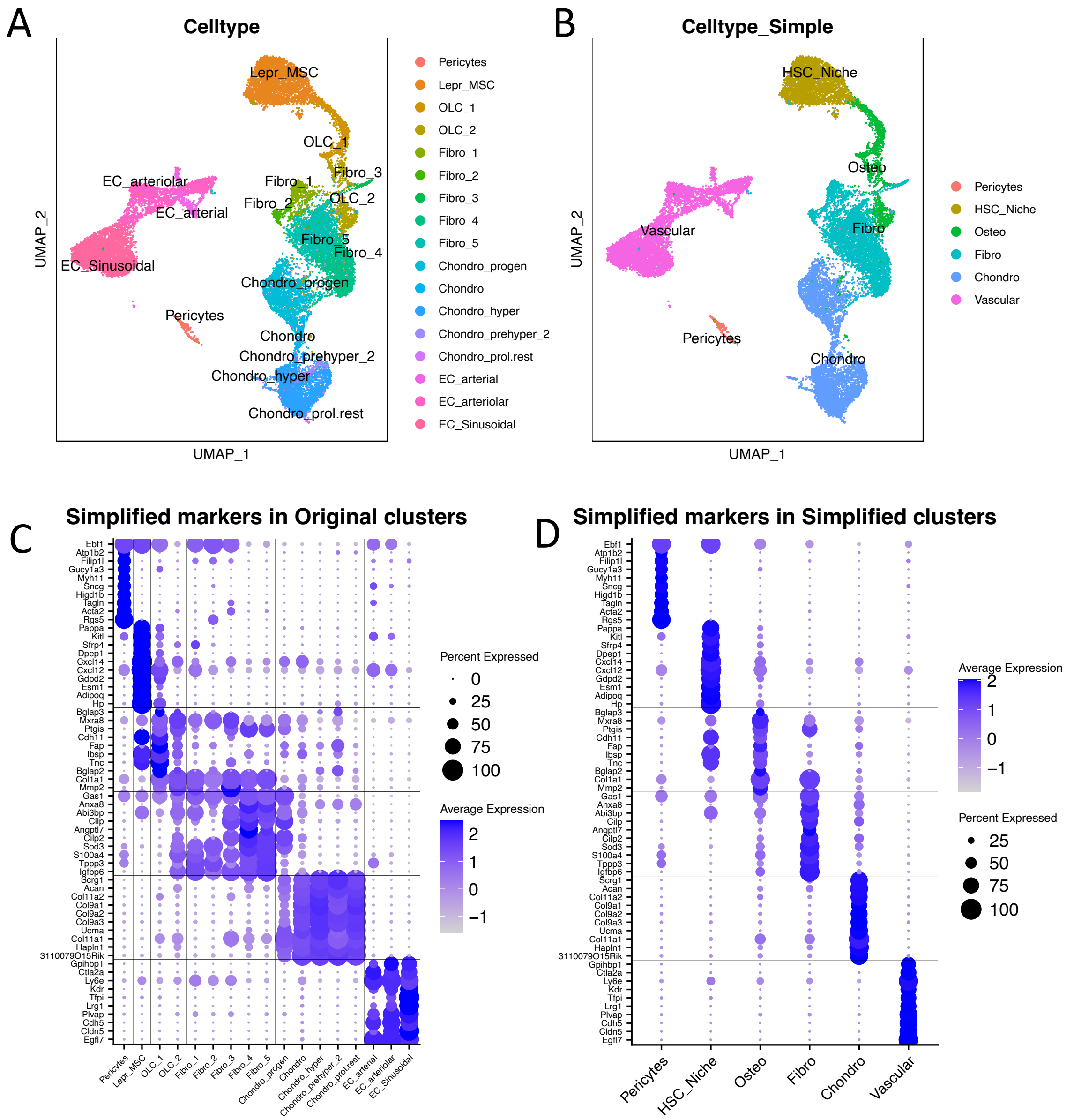

**Supplementary Figure 17. Label transfer from bone atlas to non-malignant fibroblasts and endothelial cells.** A, C: Label transfer results in OS non-malignant endothelial cells and non-malignant cancer associated fibroblasts, respectively. B,D: Label transfer scores in OS non-malignant endothelial cells and non-malignant cancer associated fibroblasts, respectively.

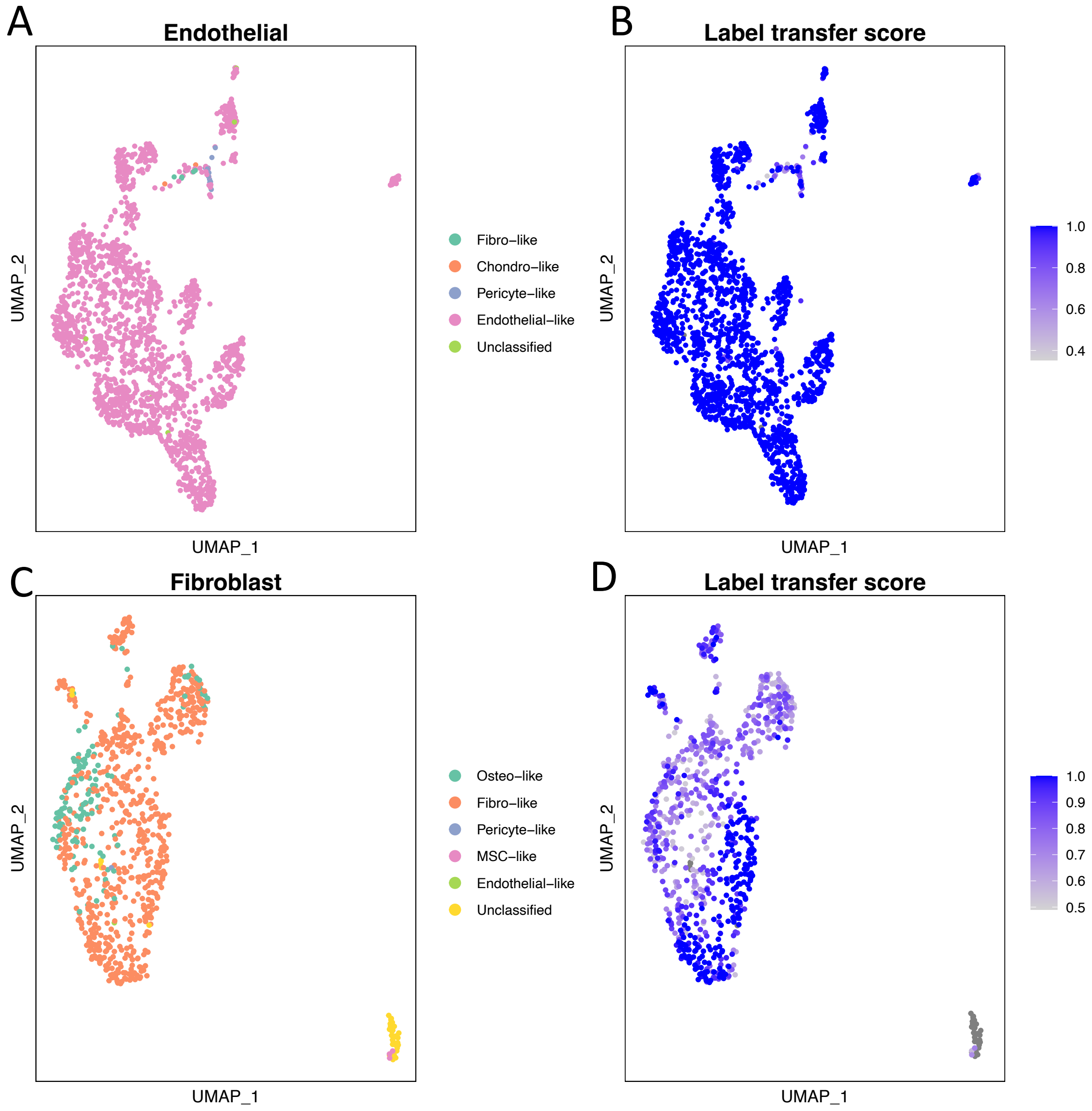

**Supplementary Figure 18. Markers and transcription factors from malignant cells stratified by pathologic subtype.** A: Dotplot showing markers from the three most common inferred celltypes among malignant cells. B: SCENIC AUCell Regulon scores of transcription factors associated with the three most common inferred celltypes among malignant cells. C: *Asb5* expression among malignant subtypes and non-malignant cells. D: *Asb5* expression among malignant subtypes and non-malignant cells, stratified by OS models.

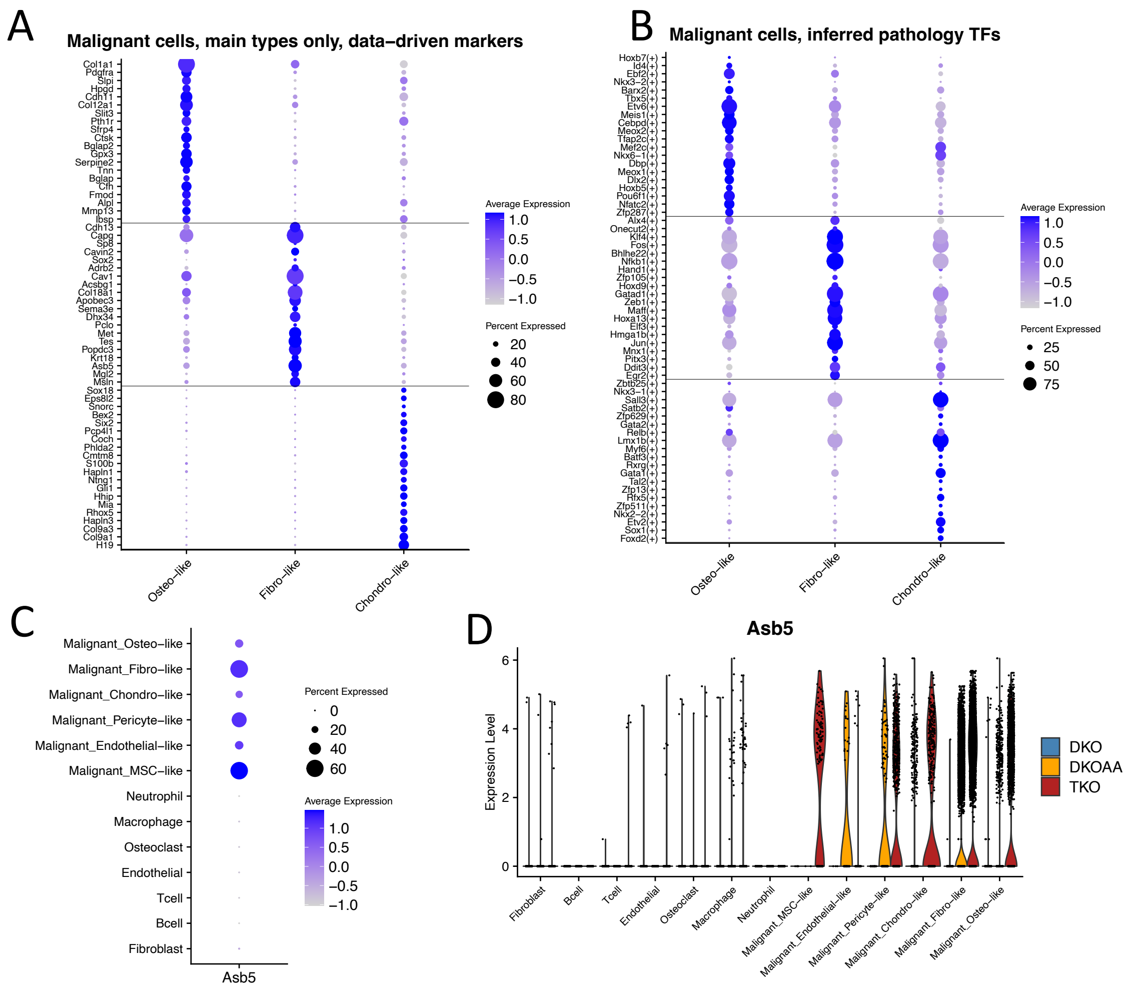

**Supplementary Figure 19. DE across OS models, stratified by pathologic subtype for which samples were available (including Osteo and Fibro-like, but not chondro-like).** A, C: TKO vs DKO differentially expressed Hallmarks gene sets in Osteo- and Fibro-like malignant cells, respectively. B,D: DKOAA vs DKO differentially expressed Hallmarks gene sets in Osteo- and Fibro-like malignant cells, respectively.

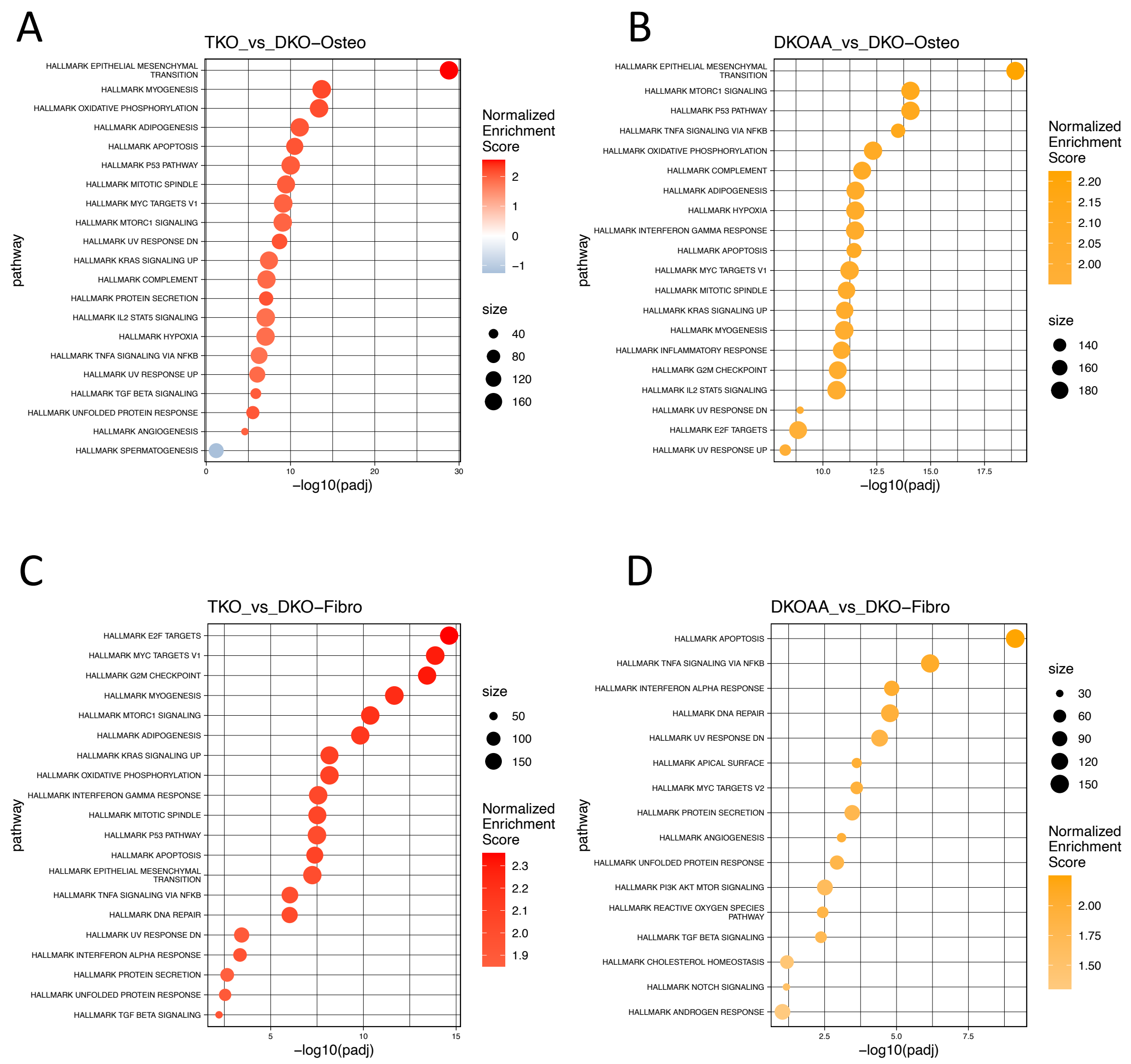

**Supplementary Figure 20. EMT gene expression in Osteo vs non-Osteo samples.** A: EMT related genes in all samples. B: EMT related genes in osteo-samples, averaged for samples in each of the three OS models. C: EMT related genes in non-osteo samples (fibro and chondro), averaged for samples in each of the three OS models.

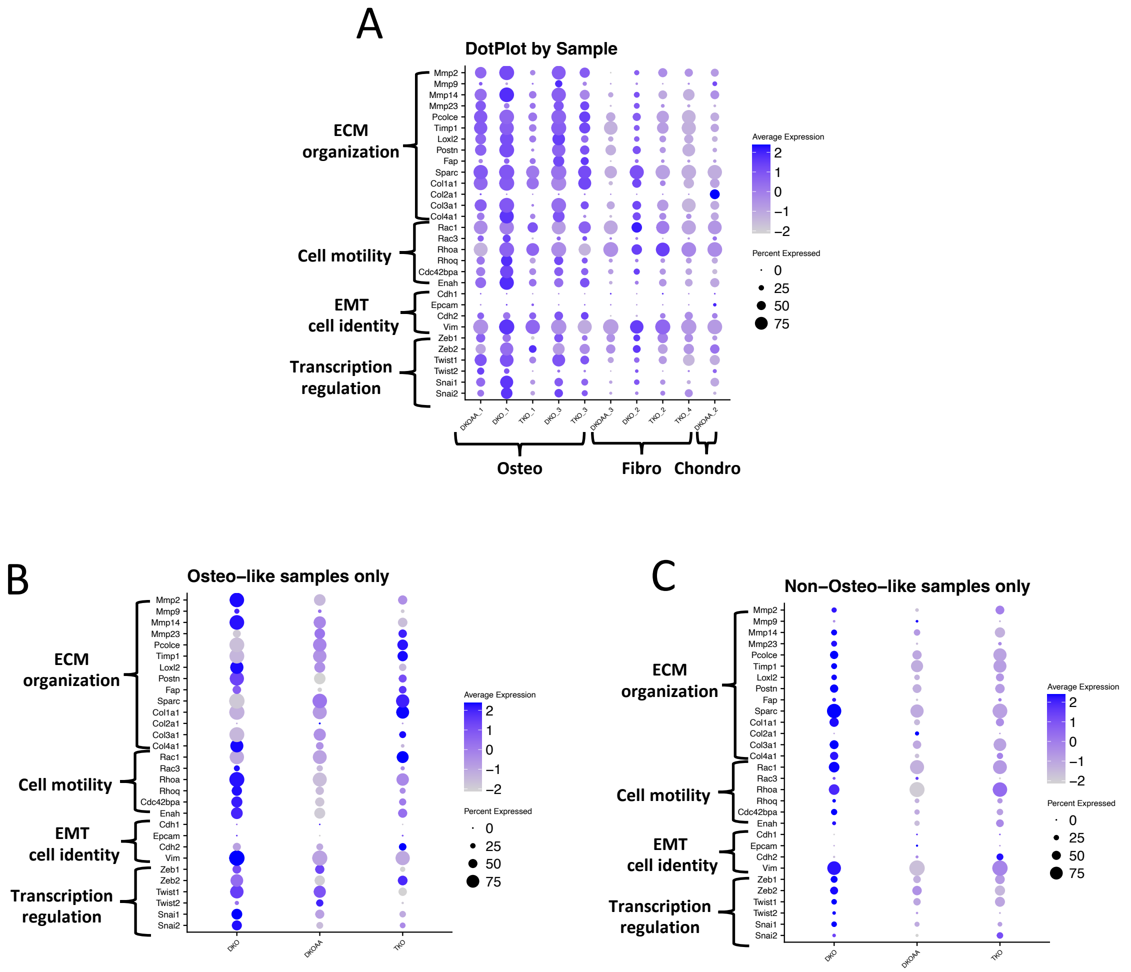

**Supplementary Figure 21. Expression of TKO myogenesis program in human OS tumors.** A: Dotplot of relevant genes, including *SKP2*, p57 (*CDKN1C*), *FBXL13*, and *ASB5*; Myogenic TFs; and the TKO upregulated Myogenesis signature. B: Featureplot showing expression score of the TKO upregulated myogenesis signature. The genes include the bottom genes of the dotplot as well as *MYOG*, the same as the signature shown in Fig 8G. C: Table showing cell numbers in clusters and patient samples. Star callout indicates the malignant cluster positive for myogenesis signal, and the associated patient sample which contributed most cells to that cluster.

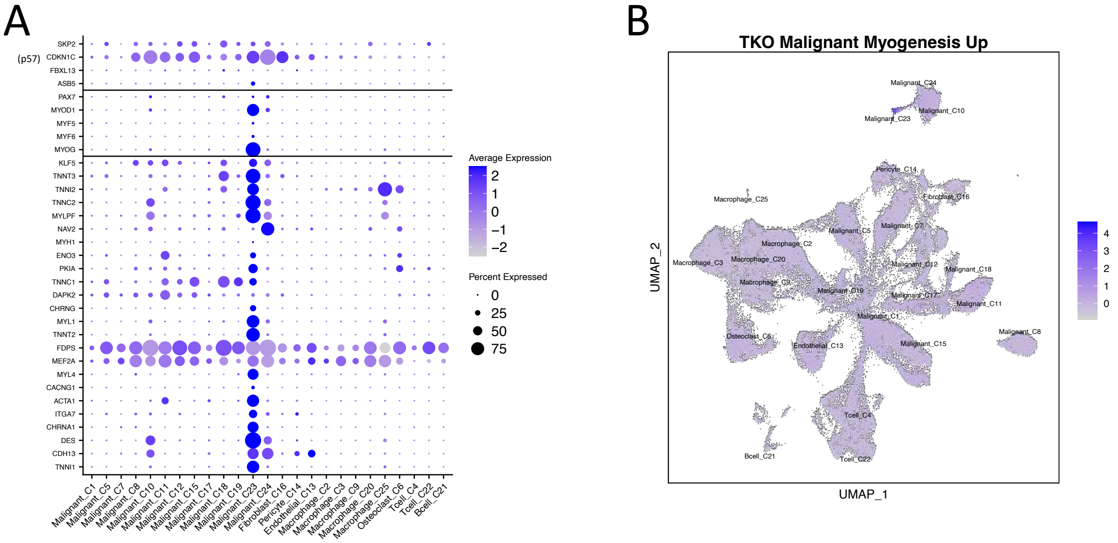


C

| Clusters | FrontOncol1_GSM4952363 | FrontOncol2_GSM4952364 | FrontOncol3_GSM4952365 | FrontOncol4_GSM5155198 | FrontOncol5_GSM5155199 | FrontOncol6_GSM5155200 | NatComm1_BC2 | NatComm2_BC3 | NatComm3_BC5 | NatComm4_BC6 | NatComm5_BC10 | NatComm6_BC11 | NatComm7_BC16 | ★ NatComm8_BC17 | NatComm9_BC20 | NatComm10_BC21 | NatComm11_BC22 |
|---|---|---|---|---|---|---|---|---|---|---|---|---|---|---|---|---|---|
| Malignant_C1 | 1048 | 417 | 203 | 60 | 55 | 351 | 361 | 980 | 1589 | 763 | 7246 | 7340 | 1424 | 450 | 346 | 1739 | 1445 |
| Malignant_C5 | 25 | 108 | 155 | 15 | 7 | 39 | 58 | 1904 | 6784 | 40 | 2765 | 225 | 129 | 13 | 5 | 24 | 73 |
| Malignant_C7 | 7 | 60 | 109 | 3 | 3 | 101 | 105 | 21 | 317 | 8095 | 178 | 10 | 11 | 13 | 26 | 72 | 16 |
| Malignant_C8 | 793 | 1556 | 5048 | 902 | 58 | 87 | 11 | 22 | 51 | 0 | 3 | 19 | 59 | 20 | 15 | 19 | 9 |
| Malignant_C10 | 0 | 0 | 0 | 0 | 0 | 0 | 0 | 0 | 0 | 0 | 0 | 0 | 0 | 5866 | 0 | 1 | 9 |
| Malignant_C11 | 14 | 76 | 15 | 119 | 11 | 147 | 12 | 9 | 14 | 0 | 16 | 106 | 16 | 77 | 0 | 4680 | 284 |
| Malignant_C12 | 55 | 215 | 310 | 104 | 63 | 1822 | 13 | 482 | 1280 | 24 | 443 | 256 | 65 | 13 | 8 | 79 | 27 |
| Malignant_C15 | 1 | 5 | 3 | 4 | 4 | 9 | 20 | 12 | 10 | 0 | 7 | 2519 | 1122 | 1 | 0 | 10 | 16 |
| Malignant_C17 | 1 | 15 | 13 | 7 | 3 | 43 | 2 | 8 | 4 | 1 | 0 | 16 | 112 | 495 | 382 | 592 | 364 |
| Malignant_C18 | 0 | 2 | 0 | 0 | 6 | 3 | 0 | 1 | 1 | 0 | 2 | 9 | 6 | 22 | 1 | 2 | 2009 |
| Malignant_C19 | 90 | 77 | 4 | 4 | 11 | 9 | 1 | 0 | 2 | 0 | 250 | 23 | 559 | 2 | 2 | 12 | 8 |
| ★ Malignant_C23 | 6 | 0 | 1 | 1 | 0 | 0 | 0 | 0 | 4 | 0 | 0 | 0 | 0 | 324 | 0 | 0 | 37 |
| Malignant_C24 | 0 | 0 | 0 | 0 | 0 | 0 | 0 | 0 | 0 | 0 | 0 | 0 | 0 | 242 | 0 | 0 | 0 |
| Fibroblast_C16 | 7 | 9 | 40 | 50 | 0 | 292 | 0 | 0 | 175 | 3 | 298 | 15 | 3 | 59 | 630 | 6 | 896 |
| Pericyte_C14 | 24 | 125 | 253 | 69 | 49 | 89 | 1552 | 157 | 409 | 167 | 143 | 29 | 355 | 38 | 63 | 797 | 59 |
| Endothelial_C13 | 38 | 232 | 184 | 78 | 141 | 143 | 674 | 273 | 721 | 387 | 267 | 1 | 407 | 218 | 370 | 1 | 285 |
| Macrophage_C2 | 139 | 272 | 63 | 42 | 159 | 51 | 1133 | 3776 | 3378 | 2842 | 305 | 223 | 1904 | 63 | 44 | 37 | 142 |
| Macrophage_C3 | 2909 | 1637 | 47 | 785 | 5334 | 250 | 23 | 46 | 12 | 84 | 2 | 10 | 595 | 796 | 342 | 474 | 593 |
| Macrophage_C9 | 309 | 307 | 53 | 752 | 294 | 797 | 519 | 168 | 169 | 630 | 406 | 80 | 623 | 284 | 356 | 70 | 453 |
| Macrophage_C20 | 280 | 48 | 0 | 16 | 180 | 7 | 1 | 2 | 0 | 13 | 1 | 1 | 101 | 60 | 13 | 23 | 28 |
| Macrophage_C25 | 0 | 0 | 0 | 0 | 0 | 0 | 0 | 0 | 0 | 0 | 0 | 0 | 0 | 143 | 0 | 0 | 0 |
| Osteoclast_C6 | 1118 | 844 | 484 | 135 | 112 | 38 | 250 | 1233 | 722 | 2062 | 148 | 0 | 582 | 124 | 1631 | 0 | 1 |
| Tcell_C4 | 287 | 689 | 68 | 462 | 1418 | 2667 | 217 | 3120 | 50 | 95 | 527 | 163 | 885 | 439 | 1114 | 0 | 781 |
| Tcell_C22 | 17 | 29 | 2 | 6 | 225 | 103 | 0 | 37 | 0 | 0 | 0 | 1 | 22 | 15 | 1 | 1 | 8 |
| Bcell_C21 | 1 | 2 | 0 | 2 | 1 | 703 | 0 | 0 | 0 | 0 | 0 | 3 | 1 | 1 | 3 | 0 | 3 |

Samples
